# Autophagy activation, lipotoxicity and lysosomal membrane permeabilization synergize to promote pimozide- and loperamide-induced glioma cell death

**DOI:** 10.1101/2020.09.23.309617

**Authors:** Nina Meyer, Lisa Henkel, Benedikt Linder, Svenja Zielke, Georg Tascher, Sandra Trautmann, Gerd Geisslinger, Christian Münch, Simone Fulda, Irmgard Tegeder, Donat Kögel

## Abstract

Increasing evidence suggests that induction of lethal autophagy carries potential significance for the treatment of glioblastoma (GBM). In continuation of previous work, we demonstrate that pimozide and loperamide trigger an ATG5- and ATG7-dependent type of cell death that is significantly inhibited with the cathepsin inhibitors E64D/Pepstatin A and the lipid ROS scavenger α-tocopherol in MZ-54 GBM cells. Global proteomic analysis after treatment with both drugs also revealed an increase of proteins related to lipid and cholesterol metabolic processes. These changes were accompanied by AKT1 (AKT serine/threonine kinase 1) inhibition and a massive accumulation of cholesterol and other lipids in the lysosomal compartment, indicative of impaired lipid transport/degradation. In line with these observations, pimozide and loperamide treatment were associated with a pronounced increase of bioactive sphingolipids including ceramides, glucosylceramides and sphingoid bases measured by targeted lipidomic analysis. Furthermore, pimozide and loperamide inhibited the activity of acid sphingomyelinase (ASM), increased lipid-ROS levels and promoted induction of lysosomal membrane permeabilization (LMP), as well as release of cathepsin B into the cytosol in MZ-54 wt cells. While LMP and cell death were significantly attenuated in ATG5/7 KO cells, both events were enhanced by depletion of the lysophagy receptor VCP (valosin containing protein), supporting a pro-survival function of lysophagy under these conditions. Collectively, our data suggest that pimozide and loperamide-driven autophagy and lipotoxicity synergize to induce LMP and lysosomal cell death. The results also support the notion that simultaneous overactivation of autophagy and induction of LMP represents a promising approach for the treatment of GBM.

## Introduction

Autophagy is an important, cellular waste disposal mechanism that encloses old and damaged lipids, proteins and organelles into double-membrane vesicles (autophagosomes) and transports them to the lysosomes for recycling into basic building blocks. Under normal circumstances, autophagy has a pro-survival function that helps cells to cope with different stress conditions like nutrient deprivation or a high burden of harmful material such as damaged mitochondria [1, 2]. However, an increasing number of studies provide evidence that when massively upregulated, autophagy can also promote cellular demise, as described in lower organisms and various types of cancer [3–6].

Glioblastoma (GBM) is the most aggressive malignant primary brain tumor in adults. Even with current standard therapy including surgical resection and subsequent radiochemotherapy with temozolomide, the prognosis for patients remains devastating with a median survival of ∼ 15 months [7, 8]. Therapy sensitivity of GBM is limited by an intrinsic resistance of these tumors to apoptotic cell death [9, 10]. Interestingly, several anticancer drugs, such as AT-101, cannabinoids and the combination of imipramine/ticlopidine have been described to trigger autophagic cell death (ACD) in GBM [6,11–13]. Hence, especially in apoptosis-refractory tumors such as GBM, the induction of ACD may represent an alternative or additive strategy to bypass therapy resistance [14, 15].

The discrimination of different cell death modalities and the respective nomenclature used in this field of research is still controversial. According to recommendations of the Nomenclature Committee on Cell Death, the assignment of ‘autophagy-dependent cell death’ requires that cell death (i) relies on an enhanced autophagic flux, (ii) should functionally depend on more than one component of the autophagic machinery and (iii) does not engage other regulated cell death modalities such as apoptosis, necroptosis or ferroptosis [16]. It has been suggested that during ACD, self-digestion reaches a point beyond survival and cells literally eat themselves to death, but the exact underlying mechanisms regulating and executing ACD remain elusive [6,17,18].

In a previous study, we performed a small scale screen with the Enzo Screen-Well™ autophagy library to identify new inducers of ACD in MZ-54 GBM wild type (wt) cells compared to MZ-54 *ATG5* and *ATG7* knockout (KO) cells generated with CRISPR/Cas9 [19]. Among the most effective compounds were loperamide hydrochloride (loperamide), an opioid receptor agonist that also inhibits voltage-gated P/Q-type Ca^2+^ channels [20, 21], and pimozide, an antipsychotic agent that antagonizes D(2)/D(3) dopamine receptors and the 5-HT7 (5-hydroxytryptamine receptor 7) serotonin receptor [22, 23]. Both drugs carry potential relevance as anticancer agents, but a better understanding of their molecular effects in GBM will be crucial before further clinical development. Therefore, the major aim of this study was to investigate the underlying mechanisms of ACD induction by these compounds and to elucidate the determinants promoting the switch from pro-survival autophagy into a death-promoting pathway. Here, we demonstrate that both drugs act by synergistic over-activation of autophagy and disruption of lipid homeostasis, culminating in autophagy-dependent lysosomal membrane permeabilization and glioma cell death.

## Results

### Loperamide and pimozide induce autophagy-dependent cell death in glioblastoma cells

To validate the induction of ACD by loperamide and pimozide [19], cell death measurements were performed by comparing MZ-54 wt cells with three different CRISPR/Cas9 *ATG5* and *ATG7* KO cell lines. Knockout of *ATG5* and *ATG7* was confirmed by Western blot analysis (Fig. S1A, B). In addition, the microtubule associated protein 1 light chain 3 (MAP1LC3B) (LC3B)-II isoform was undetectable in all KO cell lines, indicating complete functional inhibition of autophagy (Fig. S1C). The combination of imipramine hydrochloride (imipramine) and ticlopidine hydrochloride (ticlopidine), which has been described to induce ACD in glioblastoma cells [13], served as positive control. In line with previous data, imipramine + ticlopidine, pimozide and loperamide induced cell death that was significantly reduced upon inhibition of autophagy in *ATG5* and *ATG7* KO cells (Fig. 1A-C) [19]. Similar results were observed in LN-229 wt and *ATG7* KO cells after pimozide or loperamide treatment (Fig. S2A, B). Re-expression of *ATG7* in MZ-54 *ATG7* KO cells reversed the death-inhibitory effect of the *ATG7* KO after imipramine + ticlopidine, pimozide and loperamide treatment (Fig. S3A, B). This further supports the notion that this type of cell death depends on the autophagic machinery.

**Figure 1:**
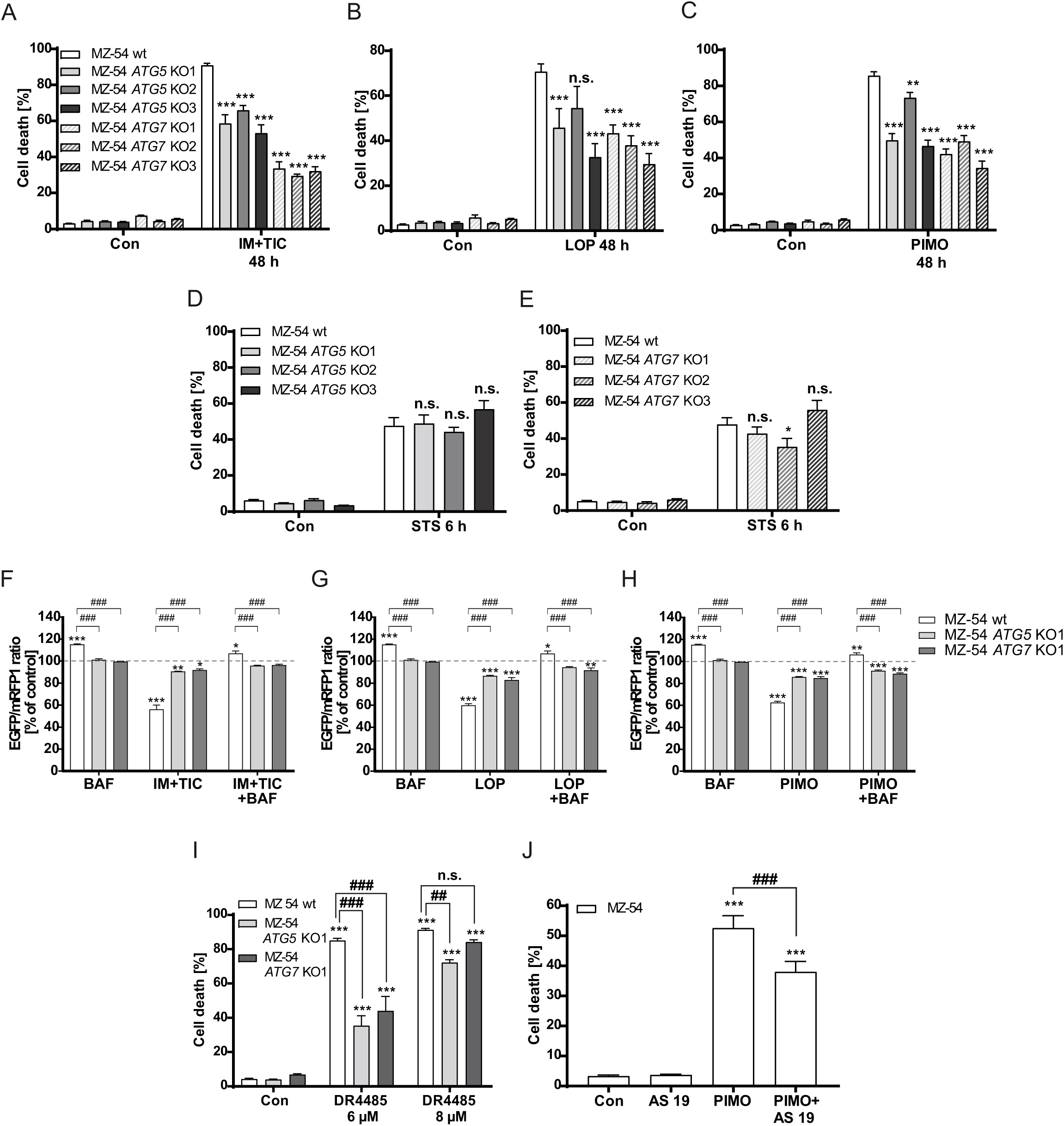
Determination of autophagic cell death upon treatment with loperamide and pimozide. (**A-E**) Flow cytometric analysis of cell death by measurement of APC-annexin V binding and PI uptake. MZ-54 wt cells and three different CRISPR/Cas9 *ATG5* and *ATG7* KO cell lines were treated with 20 µM imipramine + 100 µM ticlopidine (IM+TIC) (**A**), 12.5 µM loperamide (LOP) (**B**) 12.5 µM pimozide (PIMO) (**C**) for 48 h or with 3 µM staurosporine (STS) (**D, E**) for 6 h. **A-E** display total cell death including only-APC annexin V-positive, only-PI-positive and double-positive cells. DMSO was used as vehicle (Con, 48 h or 6 h, respectively). Data show mean + SEM of at least three experiments with three replicates and 5,000 - 10,000 cells measured in each sample. Statistical significances were calculated with a two-way ANOVA. (**F-H**) Flow cytometric analysis of the autophagic flux by measurement of the EGFP and mRFP1 signal. MZ-54 wt cells and *ATG5* and *ATG7* KO cells were exposed to 20 µM imipramine + 100 µM ticlopidine (IM+TIC) (**F**), 15 µM loperamide (LOP) (**G**) or 15 µM pimozide (PIMO) (**H**) for 16 h alone or in combination with bafilomycin A_1_ (BAF). Data show mean + SEM of three experiments with three replicates and 5,000 – 10,000 cells measured in each sample. Statistical significances were calculated with a two-way ANOVA. (**I, J**) Assessment of cell death as described above. (I) DR4485 was added in a concentration of 6 µM or 8 µM to MZ-54 wt cells or the respective *ATG5* and *ATG7* knockouts. DMSO (con) served as control. Data represent mean + SEM of three independent experiments with three replicates and 5,000 - 10,000 cells measured in each sample. Statistical significances were calculated with a two-way ANOVA. (**J**) 10 µM AS 19 was added 2 h before MZ-54 wt cells were treated with 12.5 µM pimozide (PIMO) for 40 h. DMSO (con) served as control. Data represent mean + SEM of four independent experiments with three replicates and 5,000 – 10,000 cells measured in each sample. Statistical significances were calculated with a one-way ANOVA.Significances between wt and KO cells are depicted as P ≤ 0.05: #, P ≤ 0.01: ## or P ≤ 0.001: ###.

To exclude the possibility that other mechanisms also using the LC3 conjugation machinery, in particular LC3-associated phagocytosis (LAP) [24] are responsible for the observed changes in cell death, different MZ-54 knockdown (KD) cell lines were used. The imipramine + ticlopidine-, pimozide- and loperamide-induced cell death was strongly reduced compared to the control cells when autophagy-specific genes were knocked down (*RB1CC1* and *ATG14*), but not when a LAP-specific gene (*NOX2*) was depleted (Fig. S3C-H). To rule out a general cell death resistance of *ATG5* and *ATG7* KO lines, cells were treated with the common apoptosis inducer staurosporine that elicited very similar cell death responses in wt cells and *ATG5* or *ATG7* KO cells [19] (Fig. 1D, E). Additionally, induction of the autophagic flux by loperamide, pimozide and imipramine + ticlopidine was verified by using MZ-54 wt, *ATG5* and *ATG7* KO cells stably transfected with the reporter construct pMRX-IP-GFP-LC3B-RFP-LC3BΔG. FACS analysis revealed a robust induction of the MAP1LC3B switch from MAP1LC3B-I to MAP1LC3B-II shown by a decrease in the EGFP/mRFP1 ratio that was abrogated by genetic depletion of *ATG5* or *ATG7* and by the addition of the late stage autophagy inhibitor bafilomycin A_1_ (Fig. 1F-H, see also Fig. S4A). Next, we analyzed the possible receptor dependency of loperamide- and pimozide-induced ACD. Interestingly, the serotonin receptor 5-HT7 is commonly overexpressed in malignant glioma cells and pimozide has been described to inhibit 5-HT7 receptor signaling with high affinity [25]. The potential relevance of 5-HT7 receptor signaling for ACD was determined by using the highly selective 5-HT7 receptor antagonist DR4485. Indeed, similar to pimozide, DR4485 induced autophagy as well as cell death that was diminished by knockout of *ATG5* and *ATG7* (Fig. S4B, C and 1I). Furthermore, the selective 5-HT7 agonist AS 19 could partially rescue pimozide-induced cell death, suggesting a putative role of 5-HT7 signaling in pimozide-induced ACD (Fig. 1J). In contrast to pimozide, the ACD-promoting effects of the opioid receptor agonist and voltage-gated P/Q-type Ca^2+^ channel inhibitor loperamide appear to be independent of its major receptor targets, because neither treatment with the mu-opioid receptor agonist endomorphin-1, nor inhibition of the voltage-dependent calcium channel with verapamil were able to induce cell death in MZ-54 cells (data not shown).

### Loperamide and pimozide induce upregulation of proteins involved in lipid metabolism

To investigate changes in the protein expression profile of MZ-54 cells upon exposure to loperamide and pimozide, proteomes were analyzed by a Tandem-Mass Tag -approach. A total of 6298 proteins were quantified in all samples, of which 204 and 214 proteins were significantly increased and 146 and 142 proteins were significantly reduced following loperamide and pimozide treatment, respectively (Fig. 2A and B). A list displaying all significantly changed proteins is presented in Table S1. Among the significantly upregulated GO terms we found autophagy- and lysosome-related protein clusters, which is in accordance to the increased autophagic flux induced by loperamide and pimozide. Another strongly upregulated GO term was “vesicle-mediated transport”, linking autophagy with dysregulations of lipid transport and metabolism [26–28]. Among the downregulated protein clusters, relevant GO terms were “mitotic cell cycle” (GO:0000278) as well as nucleic acid metabolic processes, pointing to growth inhibition by the compounds.

**Figure 2:**
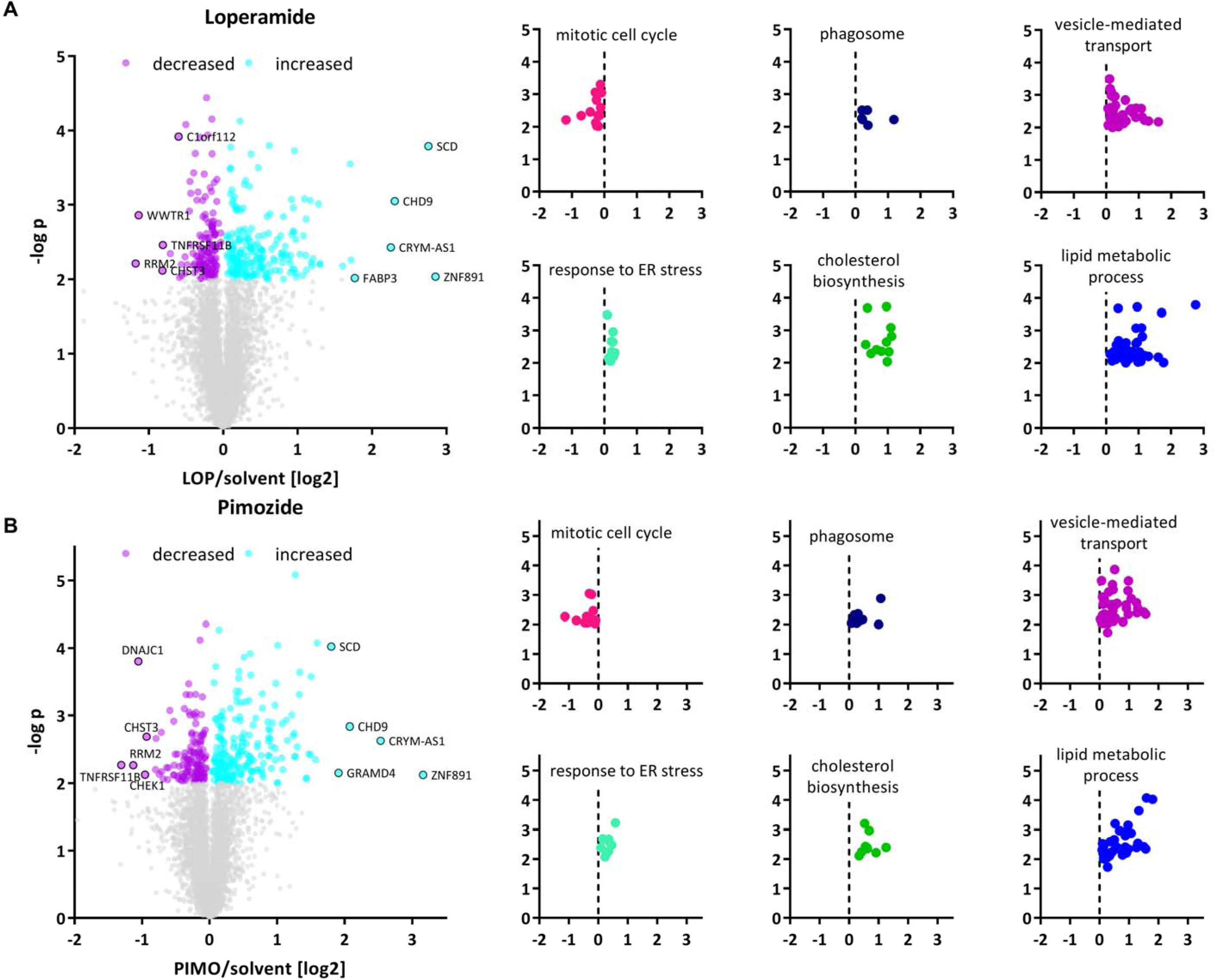
Whole proteome analysis of loperamide- and pimozide- treated MZ-54 glioma cells. (**A and B, left panel**) Volcano plots displaying the protein ratios (log2) of TMT-labeled proteome data as a function of the -log p-value. MZ-54 cells were exposed to 12.5 µM loperamide (LOP; **A**) or 12.5 µM pimozide (PIMO; **B**) for 24 h. Experiments were performed in duplicates and a total of 6298 proteins were quantified in all samples. Significantly up-or downregulated proteins with p<0.01 are each depicted in purple or light blue, proteins with no significant expression changes are shown in grey. (**A and B, right panel**) Bioinformatic analyses using String [29] revealed that significant reduction of the GO-term “mitotic cell cycle” (GO:0000278; upper left; pink) and a significant enrichment of the GO-terms “response to ER-stress” (GO:0034976; lower left; turquoise), “vesicle-mediated transport” (GO:0016192; upper right; purple) and “lipid-metabolic process” (GO:0006629; lower right; blue). Additionally, the KEGG-pathway “phagosome” (hsa04145; upper middle; dark blue) and the reactome-pathway “cholesterol biosynthesis (HSA-191273; lower middle; green) were significantly enriched after both treatments.

Remarkably, String analyses [29] revealed that treatment with both compounds led to a strong and robust increase of the gene ontology (GO) term for “lipid metabolic processes” (GO:0006629) and the reactome pathway “cholesterol biosynthesis” (HSA-191273). In brief, out of the top 50 significantly upregulated proteins by loperamide treatment, 13 (26%) were related to cholesterol metabolism and 18 (36%) were associated with lipid metabolic processes. Similar effects were obtained for pimozide-treated cells, with 9 proteins (18%) out of the top 50 upregulated proteins being involved in cholesterol metabolism and 14 (28%) being related to lipid metabolic processes. Interestingly, treatment with loperamide or pimozide induced an upregulation of the proteins SCARB1 (scavenger receptor class B member 1), APOB (apolipoprotein B), LDLR (low density lipoprotein receptor), HMGCR (3-hydroxy-3-methylglutaryl-CoA reductase), APP (amyloid beta precursor protein), APLP2 (amyloid beta precursor like protein 2), FDFT1 (farnesyl-diphosphate farnesyltransferase 1) and HMGCS1 (3-hydroxy-3-methylglutaryl-CoA synthase 1), all belonging to the GO term “cholesterol metabolic processes”. These findings indicate that treatment with these compounds strongly affect lipid turnover and/or trafficking, specifically involving cholesterol.

### Loperamide and pimozide impair lipid trafficking

It has been previously described that upregulation of proteins involved in cholesterol metabolism may be caused by impaired lipid trafficking, because decreased cholesterol levels in the cytosol promote the constitutive activation of the transcription factor sterol regulatory element binding proteins (SREBP) [30]. Of note, four of the proteins upregulated by loperamide and pimozide that were related to cholesterol metabolic processes, namely HMGCR (3-hydroxy-3-methylglutaryl-CoA reductase), LDLR (low density lipoprotein receptor), HMGSC1 (3-hydroxy-3-methylglutaryl-CoA synthase 1) and FDFT1 (farnesyl-diphosphate farnesyltransferase 1), are directly regulated by SREBP2 (sterol regulatory element binding protein 2) [31]. In line with these observations, several lipid storage diseases are associated with high expression of proteins involved in cholesterol metabolism. These diseases, e.g. Niemann-Pick Disease Type A-C or Gaucher disease, are characterized by the accumulation of lipids in lysosomes and late endosomes owing to mutations of lysosomal enzymes involved in lipid trafficking or catabolism [30,32,33].

To elucidate our hypothesis that cholesterol transport was altered upon treatment with the autophagy-inducing drugs loperamide, pimozide and imipramine + ticlopidine, cholesterol was stained with filipin III in combination with the lysosomal marker LAMP1 (lysosomal associated membrane protein 1) (Fig. 3A). While filipin III staining of vehicle-treated cells was weak and evenly distributed throughout the cells, exposure to the cholesterol synthesis inhibitor U18666A, loperamide and pimozide resulted in a strong colocalization of filipin III puncta with anti-LAMP1, indicating a massive accumulation of cholesterol in the lysosomes. The steroidal, cationic amphiphile U18666A has been previously shown to inhibit intracellular cholesterol trafficking out of the lysosomal/endosomal compartment, suggesting that impaired cholesterol transport contributes to the phenotype of loperamide- and pimozide-treated cells [34].

**Figure 3:**
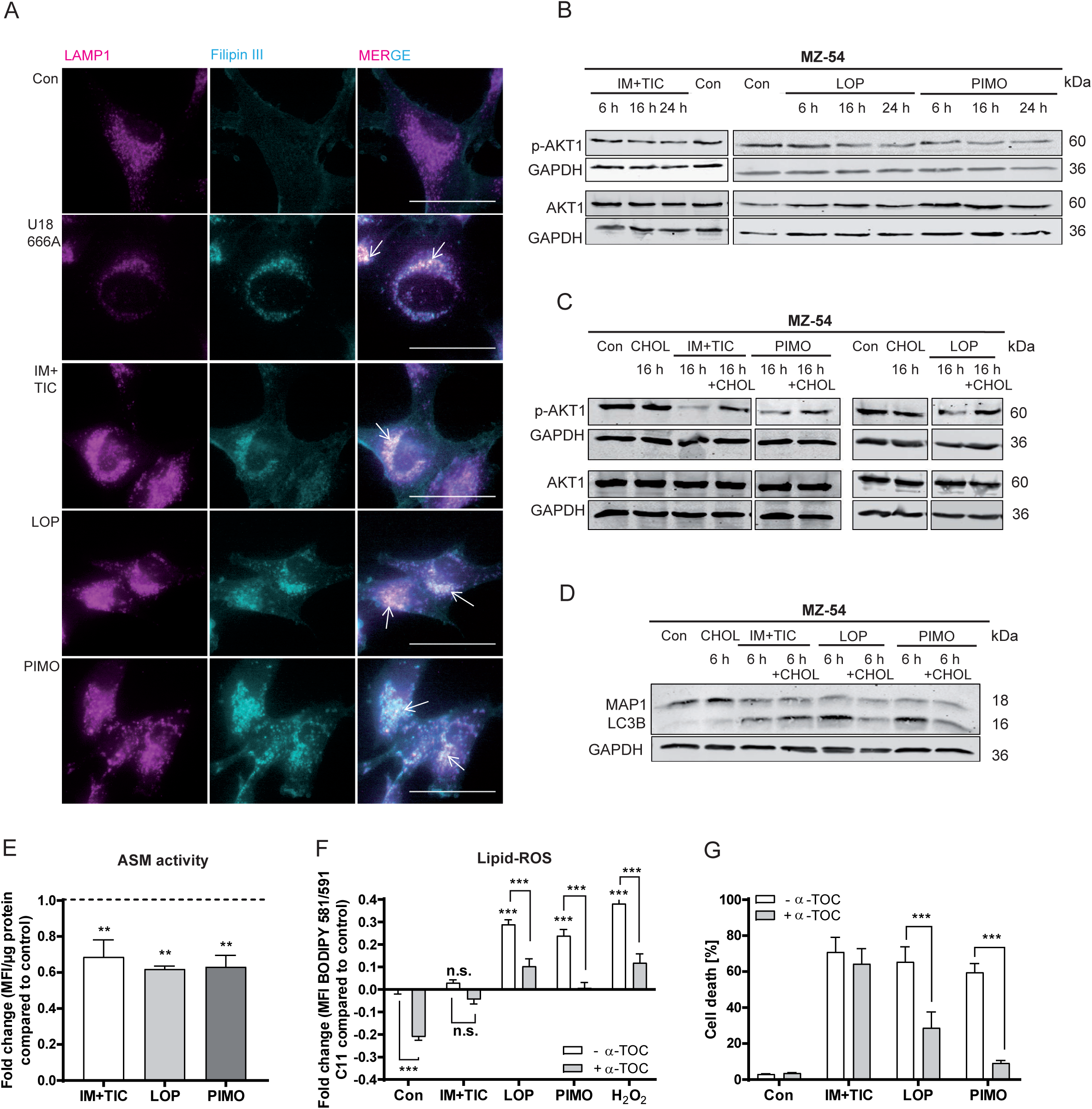
Determination of altered lipid trafficking in imipramine + ticlopidine, loperamide and pimozide treated MZ-54 cells. (**A**) Microscopic analysis of cholesterol accumulation in the lysosomes. Cholesterol was marked with filipin III and lysosomes were stained with anti-LAMP1. MZ-54 cells were treated with 1.25 µM U18666A as positive control, 20 µM imipramine + 75 µM ticlopidine (IM+TIC), 12.5 µM loperamide (LOP), 12.5 µM pimozide (PIMO) or DMSO (Con) for 18 h. At least three images were taken per experiment at 60x magnification and three independent experiments were performed (Scale bar = 50 µm). Arrows display colocalization of filipin III and LAMP1. (**B-D**) Immunoblot analysis of phospho-AKT1 and AKT1 protein expression (**B, C**) and of MAP1LC3B switch (**D**). GAPDH was used as housekeeper. MZ-54 cells were treated with 20 µM imipramine + 100 µM ticlopidine (IM+TIC), 15 µM loperamide (LOP) or 15 µM pimozide (PIMO) for the indicated time periods (6 h, 16 h, 24 h) alone or in combination with 10 µg/mL cholesterol– methyl-β-cyclodextrin. DMSO was used as control (Con). Experiments were performed two to three times. (**E**) Assessment of ASM activity upon treatment with 20 µM imipramine + 75 µM ticlopidine (IM+TIC), 12.5 µM loperamide (LOP) or 12.5 µM pimozide (PIMO) for 16 h. The bar chart represents the fold change of mean fluorescent intensity (MFI) per µg protein compared to control. (**F**) Assessment of lipid-ROS levels by flow cytometric analysis of MZ-54 cells stained with the lipid peroxidation sensor BODIPY™ 581/591 C11. Cell were exposed to 15 µM pimozide (PIMO), 15 µM loperamide (LOP), 20 µM imipramine + 100 µM ticlopidine (IM+TIC), 100 µM H_2_O_2_ (positive control) or DMSO (Con) for 16 h alone or in combination with 100 µM α-tocopherol (α-TOC, added to the cells 1 h before the other treatments). Data represent the fold change of BODIPY™ 581/591 C11 MFI compared to control. (**G**) Quantification of cell death by flow cytometric analysis of APC-annexin V binding and PI uptake. MZ-54 cells were treated with 20 µM imipramine + 75 µM ticlopidine (IM+TIC), 12.5 µM loperamide (LOP), 12.5 µM pimozide (PIMO) or DMSO (Con) for 48 h in the presence or absence of 100 µM α-tocopherol (α-TOC, added to the cells 1 h before the other treatments). Total cell death was calculated by summing up only-APC-annexin V-positive, only-PI-positive and double-positive cells. **E**, **F** and **G** represent mean + SEM of at least three independent experiments performed in triplicates (5,000 – 10,000 cells measured in each sample). Statistical significances were calculated with a two-way ANOVA.

Interestingly, increasing amounts of cholesterol in the cytosol has been reported to negatively regulate autophagy via mTOR (mechanistic target of rapamycin kinase) and AKT1 (AKT serine/threonine kinase 1) signaling. Conversely, restricting the availability of cytosolic cholesterol by impairing its transport is accompanied with the induction of autophagy [35, 36]. We found that imipramine + ticlopidine, loperamide and pimozide effectively decreased mTOR (mechanistic target of rapamycin kinase) signaling [19],AKT1 and ribosomal protein S6 kinase signaling, shown by a decrease of phosphorylated AKT1 at Ser473 and phosphorylated RPS6 at Ser240/244 (Fig. 3B and Fig. S5A). In accordance with these observations, addition of water-soluble cholesterol (cholesterol–methyl-β-cyclodextrin) increased phospho-AKT1 and phospho-RPS6 levels after treatment with all three compounds (Fig. 3C and Fig. S5B) and suppressed the MAP1LC3B switch upon exposure to loperamide and pimozide (Fig. 3D). Based on these findings we conclude that decreased cytosolic cholesterol might promote the autophagy-inducing effect of loperamide and pimozide in MZ-54 glioma cells.

Of note, previous screens by Kornhuber et al. [37, 38] revealed that imipramine, loperamide and pimozide are functional inhibitors of acid sphingomyelinase (ASM), a lysosomal enzyme that catalyzes the conversion of sphingomyelin to phosphorylcholine and ceramide. These compounds get trapped in the lysosome owing to their lipophilic and weakly basic properties, eventually impairing binding of ASM to the lysosomal membrane, leading to detachment of ASM and its subsequent inactivation [38, 39]. Consequently, the inactivation of ASM reduces *de novo* synthesis of ceramides. Consistent with these reports, we could confirm a significant decrease of ASM activity by loperamide, pimozide and imipramine + ticlopidine (Fig. 3E). Of note, it has been reported that accumulation of sphingomyelin due to ASM inhibition is accompanied by lysosomal damage and a late stage block in the autophagy pathway [40]. It should be pointed out that loss of ASM function leads to Niemann Pick disease, but a moderate reduction of its activity may be of therapeutic value [41].

Moreover, we hypothesized that lysosomes massively filled with lipids are prone to lysosomal membrane damage due to increased oxidative stress. Indeed, assessment of lipid-ROS levels with the lipid peroxidation sensor BODIPY^TM^ 581/591 C11 revealed that loperamide, pimozide and the positive control H_2_O_2_, but not imipramine + ticlopidine significantly increased cellular lipid-ROS levels, which was antagonized by addition of the lipid-ROS scavenger α-tocopherol (Fig. 3F). In line with these findings, α-tocopherol strongly diminished loperamide and pimozide-induced cell death, clearly pointing to an important role of lipid-ROS in loperamide and pimozide-induced cellular demise (Fig. 3G). In general, increased cellular ROS levels may be attributed to endoplasmatic reticulum (ER) stress, mitochondrial damage, or elevated expression of NAD(P)H quinone dehydrogenase oxidases. In contrast to our recent findings on the autophagy/mitophagy inducer AT-101 [42], loperamide and pimozide failed to induce mitochondrial depolarization (data not shown), suggesting that elevated ROS levels upon loperamide and pimozide treatment are not linked to gross mitochondrial damage. Of note, our proteomic analysis also did not show any signs of elevated NAD(P)H quinone dehydrogenase oxidases, but revealed a significant increase of the GO-term “response to ER-stress” (GO:0034976) (Fig. 2) following treatment with loperamide and pimozide, suggesting that increased cellular ROS levels might indeed be linked to ER-stress induced by these drugs.

### Loperamide and pimozide strongly increase cellular ceramides

The observed derangement of cholesterol pointed to perturbed lipid homeostasis as a major contributing mechanism for loperamide- and pimozide-dependent ACD. To further assess the role of lipids we focused on sphingolipids, which are crucial for lysosome biology and sensitive to lysosomal dysfunctions leading to accumulation of pathological species under stress conditions. Several studies have shown that ceramides, which belong to the group of bioactive sphingolipids, are capable of inducing autophagy as well as cell death [43–46].

Intriguingly, targeted lipid analyses of loperamide and pimozide-treated MZ-54 cells revealed a massive increase of nearly all measured ceramides and glucosylceramides, and sphingoid bases (Fig. 4A-C). The changes in lipid patterns after treatment with loperamide and pimozide were strikingly similar, except for the group of lactosylceramides, which were reduced by pimozide, but were not affected by loperamide, pointing to some drug-specific effects on galactosidase alpha. These data suggest a similar global effect of loperamide and pimozide on sphingolipid degradation, which cannot be assigned to a single enzyme. In particular, ASM inhibition does not sufficiently explain the observed lipid patterns because it should lead to a reduction of ceramide generation rather than to ceramide accumulation. As enhanced levels of sphingolipids could result from either increased *de novo* synthesis within the ER or impaired lysosomal degradation, we next assessed the effects of loperamide and pimozide on the expression of several enzymes involved in ceramide synthesis (Fig. S6A-F). Since only *CERS1* (ceramide synthase 1) expression was increased after pimozide exposure and all other enzymes were not affected by both compounds, we concluded that disruption of lysosomal degradation, but not *de novo* synthesis, is the predominant mechanism for accumulation of these lipids.

**Figure 4:**
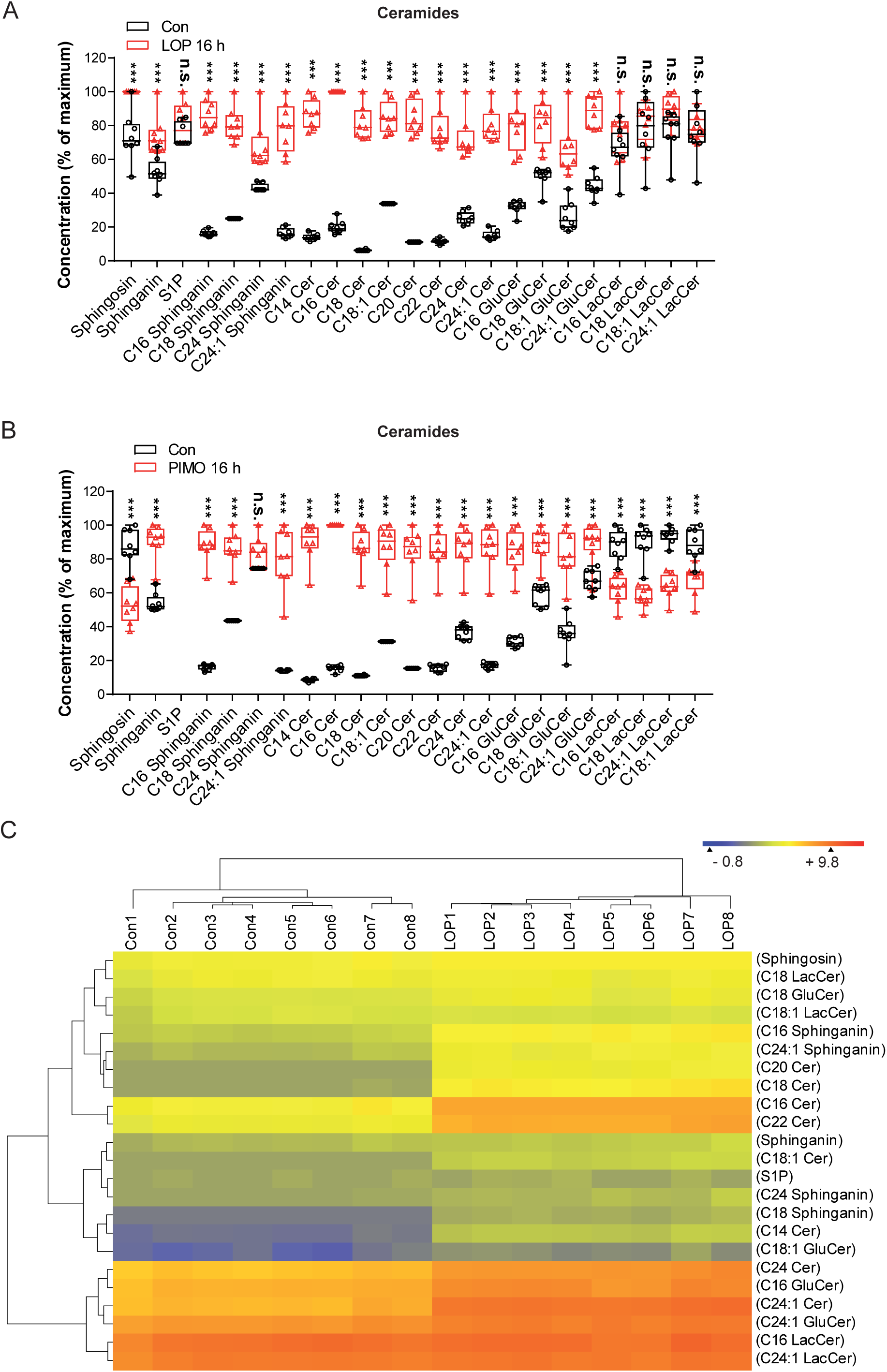
Targeted LC-MS/MS analysis of sphingolipids and ceramides in MZ-54 cells treated with loperamide or pimozide. (**A, B**) MZ-54 cells were exposed to 12.5 µM loperamide (LOP) (**A**), 12.5 µM pimozide (PIMO) (**B**) or DMSO as control (Con). FCS in the cell culture medium was reduced to 2% for loperamide and to 5% for pimozide treatments and the respective controls to minimize background. The box plots represent the interquartile range of sphingolipid- and ceramide concentrations shown as percentage of the maximum, the line is the median and whiskers show min and max values of the eight replicates, depicted as scatters. Statistical significance was assessed with a two-way ANOVA and subsequent t-tests for each lipid individually employing a correction of alpha according to Sidak. Asterisks indicate significant differences versus control, ***P<0.001. (**C**) Heatmap and dendrograms showing the result of hierarchical Euclidean clustering of lipids (y-axis) and of the individual experimental replicates (x-axis). The experimental groups are clearly separated. Colors per row display the concentration differences between different lipids and groups. The abbreviations are: S1P, sphingosine-1-phosphate; C14Cer etc. ceramide with 14 C-atoms; C24:1Cer etc., ceramide with 24 C-atoms and one unsaturated bound.

In conclusion, besides the well-proven induction of the autophagic flux and ACD by loperamide and pimozide [19], our data suggest a disruption of lysosomal sphingolipid degradation with strong accumulation of these lipids, which apparently outweighs the drug-evoked block of sphingomyelin to ceramide conversion via ASM. Hence, we propose that accumulation of ceramides and their hexosylmetabolites contributes to lysosomal dysfunction.

### Loperamide- and pimozide-induced lysosomal membrane permeabilization is strongly enhanced by ongoing autophagy

Lysosomal lipid accumulation and lipid peroxidation are both well-known inducers of lysosomal membrane permeabilization (LMP) [47]. To visualize the potential induction of LMP and to elucidate the role of autophagy in this context, MZ-54 wt cells as well as *ATG5* and *ATG7* KO cells were transfected with the pmCherry-GAL3 reporter (consisting of mCherry as fluorescent reporter and galectin-3), which translocates to the lysosomal membrane upon its rupture [48]. Interestingly, administration of loperamide and pimozide led to the formation of mCherry-GAL3 puncta that colocalized with the lysosomal marker LAMP1 in MZ-54 wt cells, clearly demonstrating the induction of LMP (Fig. 5A, B). Moreover, washout of the compounds extensively decreased the number of mCherry-GAL3 puncta per cell, suggesting an ongoing autophagic degradation of ruptured lysosomes (Fig. 5A, B). Strikingly, *ATG5* and *ATG7* KO cells showed significantly fewer mCherry-GAL3 puncta after loperamide and pimozide treatment in comparison to MZ-54 wt cells (Fig. 5C, D), thus confirming that loperamide and pimozide trigger LMP in an autophagy-dependent manner. These results could be reproduced in LN-229 wt and *ATG7* KO cells (Fig. S2C-F).

**Figure 5:**
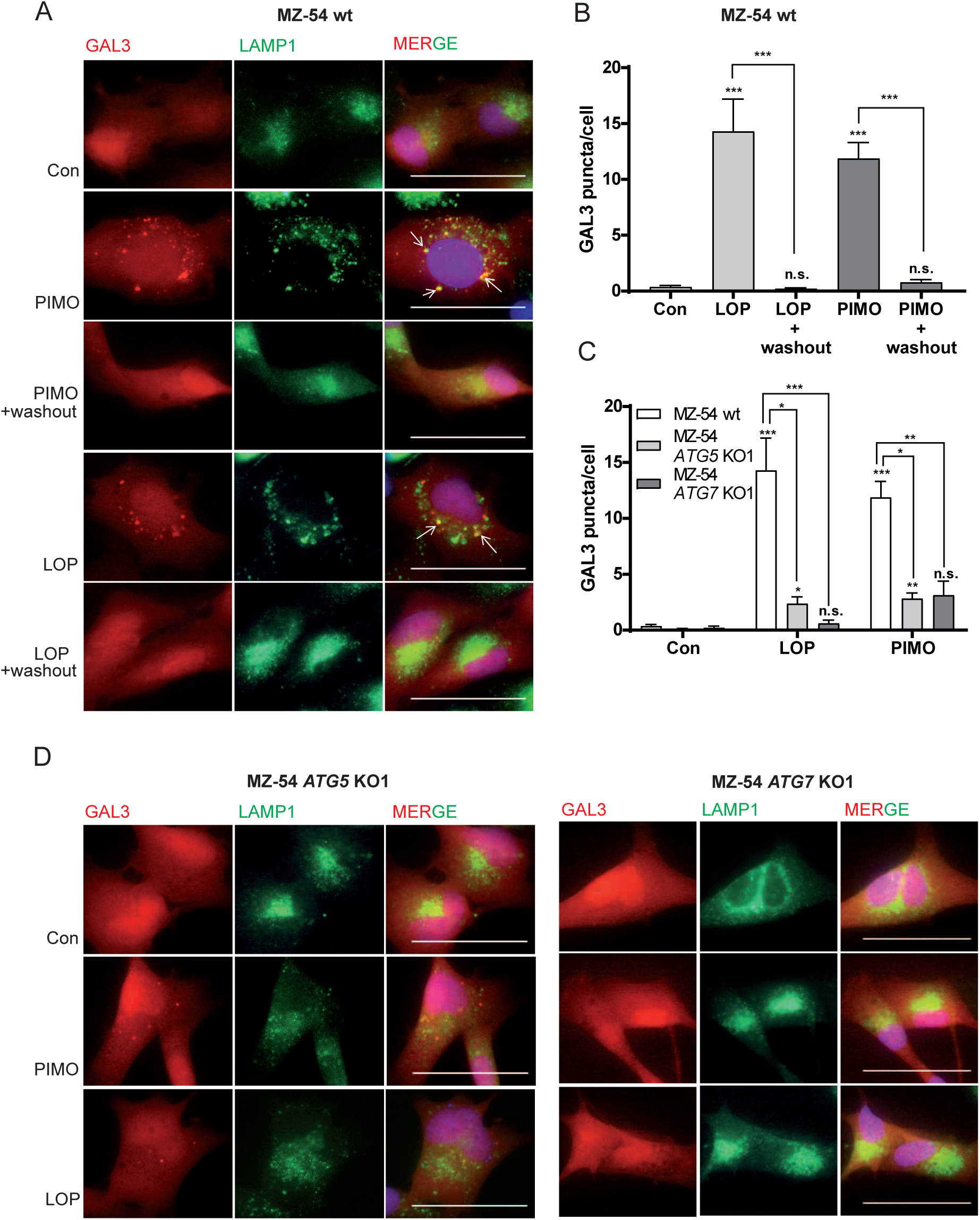
Assessment of lysosomal membrane permeabilization after loperamide and pimozide treatment. (**A-D**) Monitoring lysosomal membrane permeabilization by microscopic analysis of mCherry-GAL3 puncta formation and colocalization with the lysosomal marker LAMP1. MZ-54 control (**A**), *ATG5* and *ATG7* KO cells (**D**) stably transfected with pmCherry-GAL3 were treated with 15 µM loperamide (LOP), 12.5 µM pimozide (PIMO) or DMSO (Con) for 16 h. For washout experiments shown in **A**, loperamide and pimozide was removed after 16 h and fresh medium was added for additional 24 h. At least three independent experiments were performed and three to eight images were taken at 60x magnification in each experiment (scale bar=50 µM). (**B, C**) Quantification of mCherry-GAL3 puncta/cell. In total, 14 – 33 cells were analyzed per condition. Data represent mean + SEM of at least three independent experiments and three to eight images per experiment. Statistical significance was analyzed with a Kruskal-Wallis test.

### Loperamide and pimozide induce cathepsin release and lysosomal cell death

To validate the induction of LMP and the rescuing effects of autophagy inhibition on LMP, we next measured lysosomal cathepsin release into the cytosol by cytosolic fractionation using digitonin. Indeed, imipramine + ticlopidine, loperamide and pimozide enhanced cathepsin B activity in the cytosol, and blockage of autophagy by *ATG5* or *ATG7* KO partially attenuated these effects (Fig. 6A). In agreement with the data shown in Fig. 3, the lipid-ROS scavenger α-tocopherol was able to reduce loperamide and pimozide-driven cathepsin B release, suggesting that α-tocopherol promotes lysosomal membrane stability (Fig. 6B). Moreover, immunoblot analysis revealed a significant cathepsin B increase in the cytosolic fraction in MZ-54 wt, but not in *ATG5* KO cells (Fig. 6C-G). From these results we infer that hyperactivated autophagy by loperamide and pimozide treatment promotes lysosomal stress and LMP, which can be diminished by autophagy inhibition.

**Figure 6:**
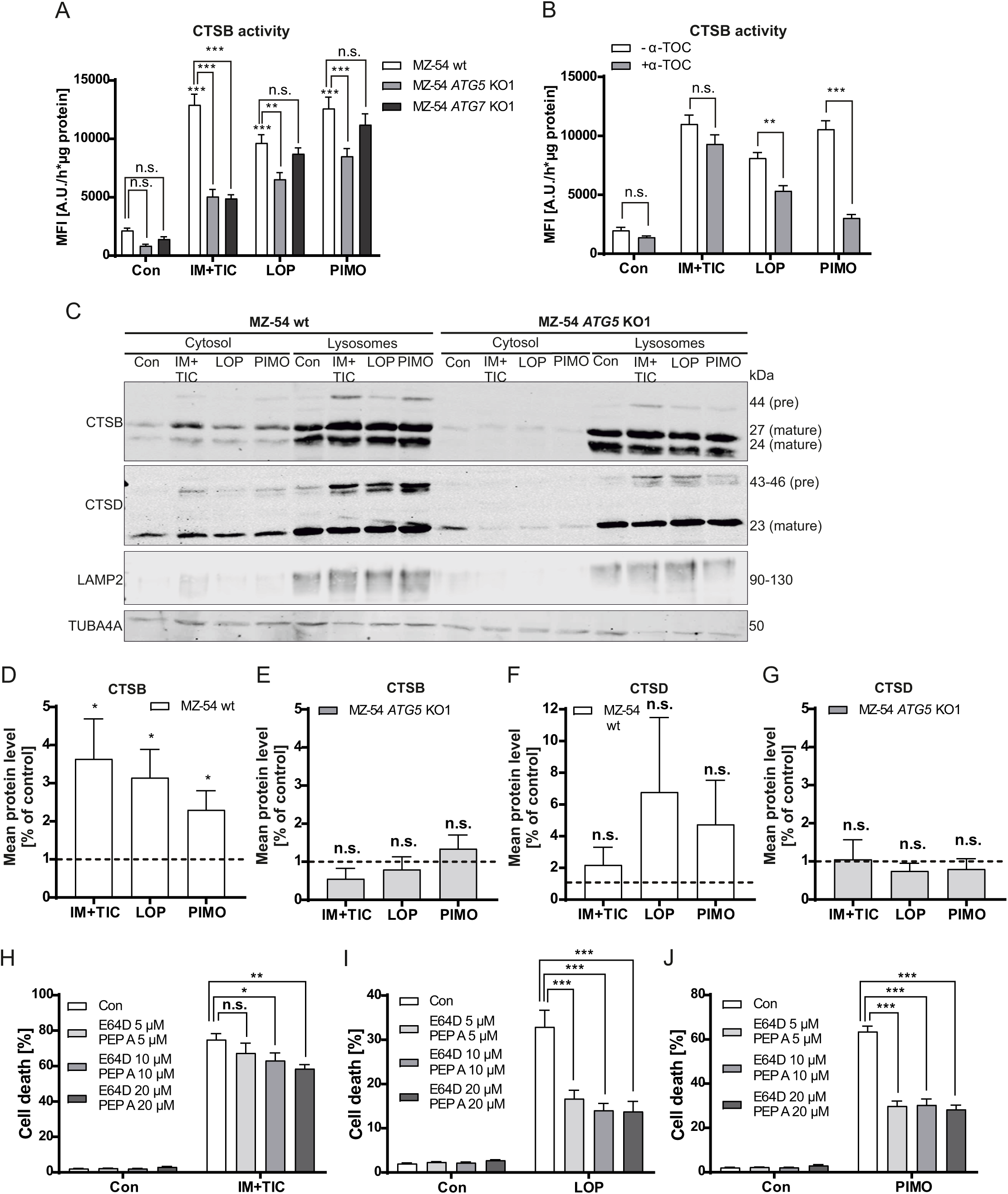
Determination of cathepsin release and lysosomal cell death. (**A, B**) Quantification of active cathepsin B in the cytosol. (**A**) MZ-54 wt, *ATG5* and *ATG7* KO cells were exposed to 20 µM imipramine + 75 µM ticlopidine (IM+TIC), 12.5 µM loperamide (LOP), 12.5 µM pimozide (PIMO) or DMSO (Con) for 16 h. Cells were fractionated using digitonin and cytosolic extracts were subjected to fluorescence-based measurement of cathepsin B activity. The bar chart represents changes in mean fluorescence intensity as arbitrary unit per h and µg protein. (**B**) MZ-54 cells were treated as in **A** in the presence or absence of 100 µM α-tocopherol (α-TOC) that was added to the cells 1 h before all other treatments. Data in **A** and **B** display mean + SEM of four independent experiments with five to six replicates. Statistical analysis was calculated with a two-way ANOVA. (**C**) Immunoblot analysis of CTSB and CTSD expression in the cytosolic fraction and organelle fraction containing lysosomes. LAMP2 was used for identification of lysosomes and TUBA4A was used for identification of cytosolic material. MZ-54 wt cells and *ATG5* KO cells were treated as described in **A**. (**D-G**) Quantification of cytosolic CTSB and CTSD protein levels of MZ54 wt cells and MZ-54 *ATG5* KO cells detected by immunoblot analysis. Data show mean + SEM of three to five independent experiments. (**H-J**) Monitoring cell death by flow cytometric quantification of APC-annexin V binding and PI uptake. Cell death refers to overall cell death including only-APC-annexin V-positive, only-PI-positive and double-positive cells. MZ-54 cells were exposed to 20 µM imipramine + 75 µM ticlopidine (IM+TIC), 12.5 µM loperamide (LOP), 12.5 µM pimozide (PIMO) or DMSO (Con) for 40 h in the presence or absence of the cathepsin inhibitors E64D and pepstatin A (PEP A) at concentrations of 5, 10 and 20 µM. Cathepsin inhibitors were added to the cells 2 h before the other treatments. Data show mean + SEM or three independent experiments with three replicates and 5,000 - 10,000 cells measured in each sample. Statistical significances were calculated with a two-way ANOVA.

It is well established that LMP has detrimental effects on cell survival, as release of the lysosomal content is accompanied with cytosolic acidification and hydrolysis of cytoplasmic material by lysosomal proteases such as cathepsins, leading to lysosomal cell death [47]. For further investigation, MZ-54 cells were exposed to loperamide and pimozide in the presence or absence of the cathepsin inhibitors E64D and pepstatin A. Indeed, cathepsin inhibition significantly attenuated the lethal effects of the drugs, indicating a relevant role of lysosomal rupture in cellular demise (Fig. 6H-J).

### Selective lysophagy induced by loperamide and pimozide promotes cell survival

Several studies demonstrated that damaged lysosomes are selectively degraded by lysophagy to circumvent the disastrous consequences of lysosomal rupture on cellular health [48–50]. Accordingly, the strong increase of mCherry-GAL3 puncta after loperamide and pimozide treatment and subsequent disappearance of mCherry-GAL3 puncta following washout of these drugs indicates ongoing lysophagy (Fig. 5A). Indeed, immunostainings revealed a clear colocalization of MAP1LC3B and mCherry-GAL3 puncta, showing the recruitment of the autophagic machinery to damaged lysosomes (Fig. 7A).

**Figure 7:**
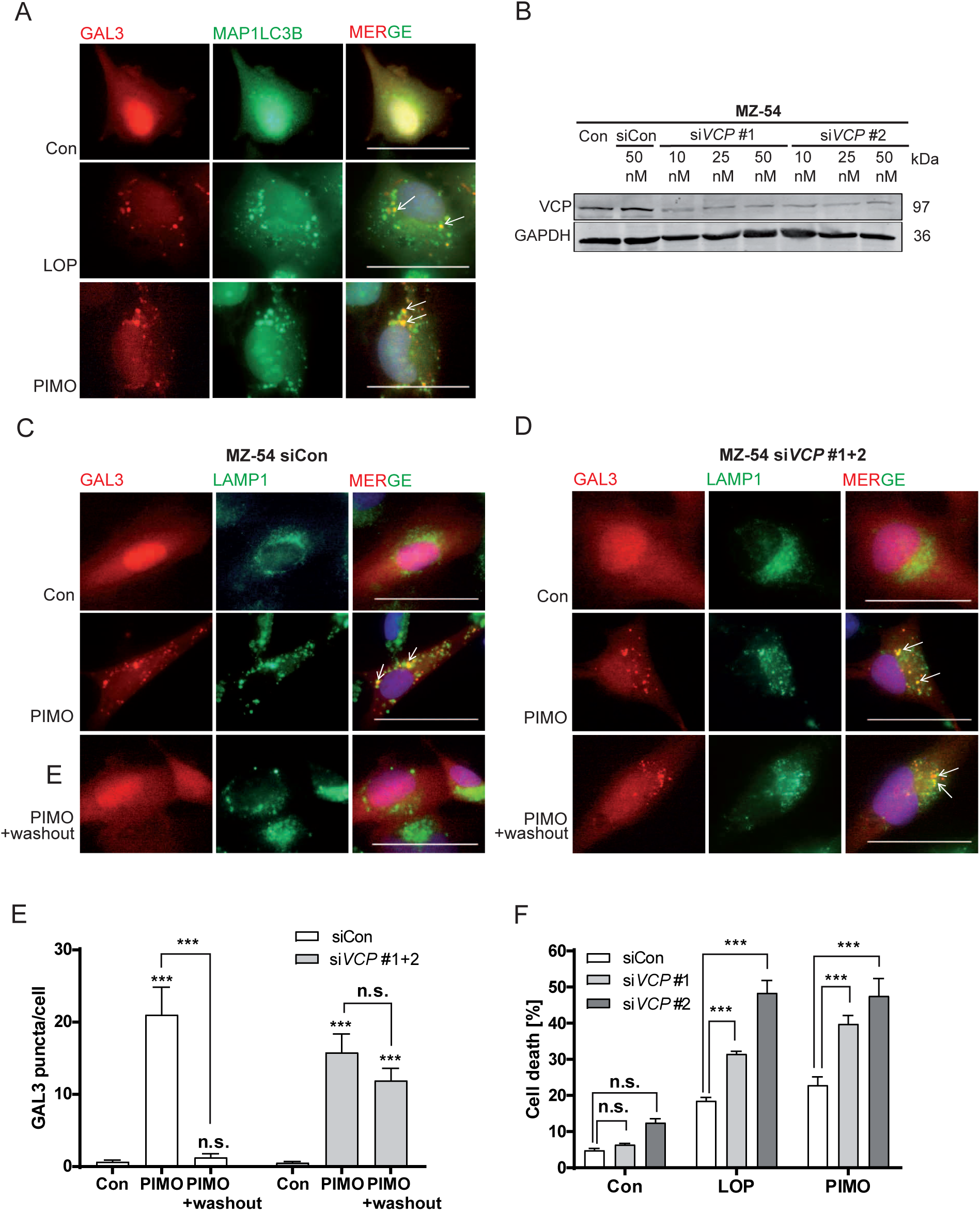
Determination of lysophagy triggered by loperamide and pimozide-induced LMP. (**A**) Microscopic analysis of mCherry-GAL3 and anti-MAP1LC3B colocalization. Images were taken at 60x magnification. Three independent experiments were performed with at least three images per experiment (scale bar = 50 µm). (**B**) Immunoblot analysis of VCP expression and GAPDH as housekeeper. MZ-54 cells were transfected with siRNA against VCP at different concentrations (si*VCP* 1 and si*VCP* 2; 10, 25 and 50 nM) or universal siRNA negative control (siCon). (**C-E**) Monitoring mCherry-GAL3 puncta formation and disappearance after compound washout by microscopic analysis of mCherry-GAL3 puncta formation and colocalization with the lysosomal marker LAMP1. MZ-54 cells were transfected with a combination of two siRNAs against VCP (25 nM si*VCP* #1 + 25 nM si*VCP* #2) or universal siRNA negative control (50 nM siCon), followed by treatment with 15 µM pimozide (PIMO) or DMSO (Con) for 16 h. For washout experiments, treatment compounds were removed after 16 h and cells were incubated in fresh medium for additional 24 h. (**C, D**) Images were taken at 60x magnification (scale bar = 50 µm). (**E**) Quantification of mCherry-GAL3 puncta per cell. In total, 11 - 23 cells were analyzed for each condition. Data represent mean + SEM of three independent experiments and three to five images per experiment. Statistical significance was analyzed with a Kruskal-Wallis test. (**F**) Flow cytometry-based cell death analyses by quantification of only-APC-annexin V-positive, only PI-positive and double-positive cells. MZ-54 cells were transfected with 25 nM siRNA against VCP (si*VCP* #1 and si*VCP* #2) or universal siRNA negative control (siCon), followed by treatment with 12.5 µM loperamide (Lop) or 12.5 µM pimozide (PIMO) for 30 h. Data represent the combined values of all three cell populations and show means + SEM of three independent experiments with three replicates and 5,000 – 10,000 cells measured in each sample. Statistical significance was calculated with a two-way ANOVA.

However, investigation of lysophagy is hitherto complicated by insufficient knowledge about the selective lysophagy receptors involved. Still, Papadopoulos et al. [48] have previously reported that the AAA+-type ATPase VCP (valosin containing protein; p97) is crucial for cell survival after lysosomal damage. Based on these findings, we assessed the putative functional role of VCP for LMP and subsequent cell death by using siRNA-mediated knockdown of VCP. Indeed, loss of VCP as confirmed by Western blot (Fig. 7B), diminished the recovery of mCherry-GAL3 puncta as marker for damaged lysosomes after pimozide washout compared to control siRNA (Fig. 7C-E). Furthermore, both siRNAs significantly increased cell death induced by loperamide and pimozide, pointing to a pro-survival function of lysophagy after induction of LMP in MZ-54 glioma cells (Fig. 7F).

## Discussion

Recently, we identified new inducers of ATG5/7-dependent cell death with potential clinical relevance for glioblastoma therapy in a mid-throughput screen of the Enzo Screen-Well™ library containing 70 known autophagy-inducing drugs [19]. For the potential clinical exploitation of ACD it is indispensable to properly decipher its underlying mechanisms that are still poorly understood. In this follow-up study, we dissected the detailed molecular events promoting activation and execution of ACD in GBM cells. To this end, we focused on two promising compounds, loperamide and pimozide, in comparison to imipramine + ticlopidine that induce ACD upon combined treatment as established in another prior study [13]. Investigation of the autophagic flux and cell death by using three different CRISPR/Cas9 KO cell lines deficient for *ATG5* or *ATG7* confirmed that all compounds clearly induced ACD in MZ-54 GBM cells. Our proteomic analysis showed an unexpected, strong increase of proteins involved in lipid and cholesterol metabolic processes. Subsequent experiments revealed that the positive association of these processes was caused by an impairment of lipid trafficking and ASM inhibition, supposed to result in accumulation of cholesterol and other lipids in the lysosomes. In line with these observations, two studies of Kornhuber et al. [37, 38] categorized imipramine, loperamide and pimozide as FIASMAs (functional inhibitors of acid sphingomyelinase), representing cationic, amphiphilic drugs that accumulate in acidic cellular compartments including lysosomes and thereby inhibit the function of ASM. The fact that both ACD-inducing compounds described in the study by Shchors et al. [13] and two out of three of our hit compounds [19] inhibit ASM suggests that ASM blockade is a common denominator and key driver in many paradigms of ACD. However, it is likely not sufficient to fully explain the observed massive increase of ceramide species upon treatment with the candidate drugs and failure of AKT1 activation in this study. The latter phenomenon is in agreement with published findings [35, 36] and with our previous studies showing that the compounds inhibit downstream mTOR (mechanistic target of rapamycin kinase) activity [19]. In general, the high cholesterol demand of several cancers including melanoma, lymphoblastoma, neuroblastoma, non-small cell lung cancer and breast adenocarcinoma has been reported to sensitize these tumors to inhibitors of cholesterol trafficking and biogenesis [51, 52]. Importantly, cholesterol metabolism in the human brain has to be tightly regulated, because the blood-brain barrier impedes uptake of lipoprotein-bound cholesterol. As such, Villa et al. [53] found that glioblastoma cells rely on the uptake of exogenous cholesterol, while normal human astrocytes are able to cover their own cholesterol demand and those of adjacent cells by *de novo* synthesis suggesting that glioblastoma cells may be particularly vulnerable to intracellular cholesterol shortage.

Besides the autophagy-inducing effect of impaired lipid trafficking, massive accumulation of lipids in the lysosome was accompanied with oxidative stress, as shown by increased lipid-peroxidation after loperamide and pimozide treatment, and subsequent induction of LMP. Interestingly, several studies reported that lysosomes massively filled with lipids become severely damaged and cannot fulfill their normal degradative and recycling properties for autophagic cargo [40,47,54,55]. Hence, in addition to accumulating waste, cells may succumb to nutrient deficiency and secondary suppression of translation and transcription, which would explain the enrichment of the related GO-terms for the downregulated proteins.

Our lipid analyses revealed a strong increase of ceramides, their glucosyl-metabolites and sphingosine/sphinganine precursors under loperamide and pimozide treatment, which was not a result of increased *de novo* synthesis, hence pointing to impaired lysosomal degradation of sphingolipids. The accumulation of bioactive sphingolipids in spite of ASM inhibition suggests that these cationic, amphiphilic drugs are also capable of inhibiting lysosomal acid ceramidase and further ceramide metabolizing enzymes, which are responsible for hydrolysis of ceramide to sphingosine [54] or degradation of glucosylceramides. Indeed, C16:0-Cer accumulation in the lysosome was found to trigger LMP by formation of ceramide channels [56]. Hence, loperamide and pimozide may have a similar effect. In addition, ASM inhibition may also trigger the increase of ceramides in the ER, thereby inducing ER stress and secondary autophagy [57].

Of note, lysosomal function is often upregulated in cancer cells to fulfill increased metabolic demands, and elevated cathepsin expression is associated with increased invasiveness and metastasis [58–61]. This high dependence on the lysosomal pathway paired with decreased stability of cancer cell lysosomes (e.g. by degradation of LAMP1 and LAMP2), as reported by several studies, makes lysosomes a particularly suitable target for cancer therapy [62, 63]. In support of this idea, loperamide and pimozide-induced LMP triggered lysosomal cell death in our cell models, which depended on the release of cathepsins into the cytosol, and was further enhanced by inhibition of lysophagy. Hence, our data support the hypothesis that lysophagy is essential for cell survival under conditions of lysosomal damage [48].

Strikingly, autophagy inhibition by functional *ATG5* and *ATG7* KO strongly decreased both LMP and cell death following drug treatment. Based on this observation, we propose that the cell-killing effects of the drugs caused by impaired lipid trafficking and lipotoxicity in the lysosomal compartment are further exacerbated by the parallel overactivation of autophagy, because lysosomal function is further challenged by the vast demand in handling the additional autophagic cargo. This demand appears to be strongly reduced in ATG KO cells as indicated by the almost complete absence of mCherry-GAL3 puncta (Fig. 5C, D), even though. lysophagy is also blocked in these cells. Collectively, our findings suggest that autophagy overactivation and lipid accumulations induced by loperamide and pimozide synergize to promote lysosomal rupture and subsequent cell death in autophagy-competent cells. Accordingly, it should be considered that ablation of autophagy (e.g. by *ATG5*/*7* KO) does not only affect ACD in general, but also prevents LMP and subsequent lysosomal cell death, at least in cases of ACD induced by ASM-inhibiting drugs. These considerations emphasize the difficulty to clearly separate autophagy-dependent cell death and lysosomal cell death, as demanded by the leading experts in the field of cell death research [16]. Nevertheless, our candidate drugs induce an ACD phenotype of cell death including increased autophagic flux, and rescue by genetic inhibition of autophagy, which is not observed with inducers of apoptosis, necroptosis and ferroptosis [19]. Our data here provide further evidence that impaired lipid trafficking promotes autophagy plus autophagy-dependent LMP and subsequent lysosomal cell death, suggesting that these mechanisms contribute to the cellular demise induced by these compounds. Moreover, our study accentuates the dual role of autophagy, showing that selective lysophagy promotes cellular survival by eliminating harmful, damaged lysosomes, while bulk autophagy rather serves to trigger cellular demise in our experimental setting. Based on the fact that (bulk) autophagy and LMP both significantly contribute to loperamide- and pimozide-induced cell death, the term ‘autophagy-dependent lysosomal cell death’ may also be used for this type of cell demise that has distinct features in comparison to other already established cell death paradigms [16].

We conclude that massive autophagy induction paired with lysosomal damage and mounting impairment of lysosomal degradative function at later time points leads to tremendous autophagic and lysosomal stress, and finally to cell death, with possible implications for GBM therapy. From a clinical standpoint, it should be considered that loperamide, an orally applied antidiarrheal agent, is not systemically bioavailable after oral administration and does not cross the blood brain barrier, although attempts have been made to target loperamide to the brain using nanoparticles [64]. In contrast, pimozide and other antipsychotic agents acting as antagonists of D(2)/D(3) dopamine receptors and/or 5-HT7 receptors [65–67] have recently gained significant attention as potential candidates for the treatment of GBM. In this context it is worth mentioning that similar to pimozide, several other antipsychotic drugs have been reported to functionally inhibit ASM, thereby affecting lipid transport [30,68,69]. Here, we provide novel evidence that pimozide-induced ACD is partly dependent on inhibition of serotonin receptor 5-HT7 signaling. Interestingly, serotonin receptor 5-HT7 is overexpressed in GBM and was previously linked to increased mitogen-activated protein kinase 1/3 (MAPK1/3) activation and interleukin 6 (IL-6) synthesis, signaling events that have been proposed to enhance glioma cell survival [23, 25]. Based on these considerations and our findings on the putative role of 5-HT7 blockade in ACD, pimozide (and other 5-HT7 receptor antagonists) might be of particular interest for glioblastoma therapy, because they are approved for clinical use and hence available for repurposing as anti-cancer agents.

## Methods

### Cell culture

DMEM GlutaMAX (10566016), penicillin/streptomycin (15140122), fetal bovine serum (FBS) (10270106), Dulbecco’s phosphate buffered saline (DPBS) (14190094) and trypsin-EDTA (25200056) were purchased from Gibco® (Life Technologies). The human glioma cell lines MZ-54 (cells were isolated from a recurrent grade IV glioblastoma [70]) and LN-229 were cultivated in DMEM, high glucose, glutaMAX, 10% fetal bovine serum and 100 U/mL penicillin-streptomycin at 37°C and 5% CO_2_. Authentication of MZ-54 and LN-229 cells was done by STR profiling at DSMZ (Sammlung von Mikroorganismen und Zellkulturen GmbH). Cells were exposed to a combination of imipramine hydrochloride (Sigma-Aldrich, 10899) and ticlopidine hydrochloride (T6654, Sigma-Aldrich) (IM+TIC), loperamide hydrochloride (LOP; Enzo Life Science, ALX-550-253), pimozide (PIMO; Sigma-Aldrich, 2062-78-4), staurosporine (STS; Enzo Life Science, 380-014-M001), AS 19 (Tocris Bioscience, 1968), DR 4485 hydrochloride (DR4485; Tocris Bioscience, 5005), cholesterol**-**methyl-β- cyclodextrin (CHOL; Sigma-Aldrich C4951), hydrogen peroxide 30% (H_2_O_2_, Merck, 1072090250) and bafilomycin A_1_ (LC Laboratories, B-1018) for the indicated time periods. MZ-54 and LN-229 cells were tested monthly for mycoplasma contamination.

### CRISPR/Cas9 knockouts

The guide RNAs for human *ATG5* and *ATG7* knockouts were selected with the online tool benchling and contained 20 nucleotides complementary to the target locus in the genome, cloning sites for BbsI and an additional 5’ *G* in case the sgRNA started with another base. Appropriate guide RNAs were separately cloned into the SpCas9(BB)-2A-GFP (PX458) vector (Addgene, 48138) or pSpCas9(BB)-2A-Puro (PX459) vector (Addgene, 48139) from Feng Zhang, respectively [71].

Both vectors encode the cas9 endonuclease from *Streptococcus pyogenes*, which requires a preceding 5’NGG protospacer adjacent motif at the genomic target locus [71]. For generation of *ATG5* and *ATG7* knockouts, a combination of two vectors containing different guide RNAs was transfected into MZ-54 cells by using Lipofectamine 3000 (Thermo Fisher Scientific, L3000008) according to the manufacturer’s instructions (DNA:Lipofectamine 3000 ratio 1:1.5). Following guide RNAs were used for the different clones: *ATG5* KO1 (*CACCGTCAGGATGAGATAACTGAAA* and *CACCGCCTCTAATGCTACCACTCAG*), *ATG5* KO2 (*CACCGAAGATGTGCTTCGAGATGTG* and *CACCGCCTCTAATGCTACCACTCAG*), *ATG5* KO3 (*CACCGTCAGGATGAGATAACTGAAA* and *CACCGCCTCTAATGCTACCACTCAG*), *ATG7* KO1 (*CACCGAATAATGGCGGCAGCTACGG* and *CACCGAAAGCTGACACTATACTGG*), *ATG7* KO2 (*CACCGAATAATGGCGGCAGCTACGG* and *CACCGAGAAGAAGCTGAACGAGTAT ATG7* KO3 (*CACCGAGAATCAGCTTGACAACAC* and *CACCGAAAGCTGACACTATACTGG*). The LN-229 ATG7 knockout line was generated with *ATG7* KO2 and *ATG7* KO3.

GFP-positive cells containing the PX458 vector were sorted into a 24-well plate 72 h after transfection by using a FACS Aria II cell sorter (BD Biosciences, Heidelberg, Germany). Cells transfected with the PX459 vector were selected with 1 ug/mL puromycin (Santa Cruz Biotechnology, sc-108071B) 48 h after transfection. After 72 h incubation, cells were separated by single cell dilution and the expanded single cell colonies containing either *ATG5* or *ATG7* KO were detected by PCR and immunoblot analysis.

### Transfection of siRNA and pmCherry-GAL3 plasmid

For depletion of VCP, MZ-54 cells were transfected with two different siRNAs against VCP (SASI_Hs01_00118726 [*VCP* #1] and SASI_Hs01_00118728 [*VCP* #2], Sigma-Aldrich) or siRNA Universal Negative Control (Sigma-Aldrich, SIC001) at 60% - 80% confluency by using Lipofectamine 3000 (Thermo Fisher Scientific, L3000008) according to the manufacturer’s manual. Therefore, the following amount of Lipofectamine 3000 was used per well: 5 µL/6-well, 1.5 µL/24-well, 1 µL/chamberslide-well. Treatments were performed 24 - 48 h after transfection. PmCherry-GAL3 was a gift from Hemmo Meyer (Addgene, 85662) [48]. Cells were transfected with pmCherry-GAL3 by using Lipofectamine 3000 according to manufacturer’s instructions. To this end, 120,000 cells were seeded into 6-well plates and cells were transfected with 2.5 µg plasmid DNA and 3.75 µL Lipofectamine 3000 (DNA: Lipofectamine ratio 1:1.5) in fresh medium on the next day. Medium was changed after 24 h and cells with stable integration of pmCherry-GAL3 were selected with 1 mg/mL G418. However, the selected cell population also contained some cells without pmCherry-GAL3 expression, which gained G418 resistance.

### Re-expression of *ATG7* in *ATG7* KO cells

For the re-expression of *ATG7* in *ATG7* KO cells, we used an inducible plasmid that only allows expression of *ATG7* in the presence of doxycycline (doxycline hyclate, D9891, Sigma Aldrich). MZ-54 *ATG7* KO3 cells were first transduced with the plasmid containing the h*ATG7* sequence (pLV[Exp]-Puro-TRE3G>hATG7[NM_001349232.1]) and in a second step transduced with the regulator plasmid (pLV[Exp]-CMV>Tet3G/Hygro) (Vectorbuilder).

Virus particles were generated in HEK 293-T cells after transfection with the above-mentioned plasmids in combination with the gag/pol plasmid psPAX2 (Addgene, 12260) and the VSV-g-envelope plasmid pMD2.G (Addgene, 12259). The plasmids were diluted in Opti-MEM and Fugene HD transfection reagent (Promega). The DNA-mixture was incubated for 30 min and then added drop-wise to the cells cultured in DMEM without penicillin/streptomycin for 6-7 h. After incubation medium was aspirated, cells were washed with DPBS and fresh culture medium was added. 24 h and 48 h after transfection the virus-containing supernatant was harvested and stored short-term at 4 °C. For transduction, 120,000 MZ-54 *ATG7* KO3 or MZ-54 *ATG7* KO3 + TRE3G-*ATG7* cells were seeded out in 6-well plates. Next day cells were incubated for 24 h with the virus-containing supernatant diluted in culture medium and 3 µg/mL Hexadimethrine bromide (polybrene, Sigma Aldrich). Afterwards medium was changed and after additional 24 h the medium was changed to selection medium containing 1 µg/mL puromycin (Puromycin dihydrochloride, sc-108071B, Santa Cruz Biotechnology) and after the second transduction also 125 µg/mL hygromycin (Hygromycin B, 10687010, Invitrogen). Re-expression was confirmed by immunoblot analysis.

### Generation of shRNA knockdowns

For depletion of *RB1CC1* virus particles were generated in HEK 293-T cells transfected with the control vector pGIPZ or the respective shRNA plasmid (KD1: RHS4430-101100096 or KD2: RHS4430-101103887) (Thermo Scientific Open Biosystems), the gag/pol plasmid pCMV-dr8.91 and the VSV-g envelope plasmid pMD2.G (Addgene, 12259). The plasmids were diluted in H_2_O and 2.5 M CaCl_2_. This DNA/CaCl_2_ solution was added drop-wise to 2xHBS solution while blowing air bubbles through a pasteur pipette and incubated for 15-20 min. Culture medium was changed and 25 µM chloroquine was added. The DNA mixture was added drop-wise to the cells and incubated for 6-8 h at 37 °C and 5 % CO_2_. After incubation the medium was replaced with fresh medium. 24 h and 48 h after transfection the virus-containing supernatant was harvested and stored short-term at 4 °C.

For depletion of *ATG14* and *NOX2* virus particles were generated in HEK 293-T cells transfected with the control vector pLKO.1 (pLKO.1 non-target) or the respective shRNA plasmid (shATG14 (KD1: TRCN0000142849 or KD2: TRCN0000144080) or shNOX2 (KD1: TRCN0000064590 or KD2: TRCN0000064591) (Sigma Aldrich), the gag/pol plasmid psPAX2 (Addgene, 12260) and the VSV-g envelope plasmid pMD2.G (Addgene, 12259). The plasmids were diluted in Opti-MEM and Fugene HD transfection reagent (Promega). The DNA-mixture was incubated for 30 min and then added drop-wise to the cells cultured in DMEM without penicillin/streptomycin for 6-7 h. After incubation medium was aspirated, cells were washed with DPBS and fresh culture medium was added. 24 h and 48 h after transfection the virus-containing supernatant was harvested and stored short-term at 4 °C.

For transduction, 120,000 parental MZ-54 cells were seeded out in 6-well plates. Next day cells were incubated for 24 h with the virus-containing supernatant diluted in culture medium and 3 µg/mL Hexadimethrine bromide (polybrene, Sigma Aldrich). Afterwards medium was changed and after additional 24 h the medium was changed to selection medium containing 1 µg/mL puromycin. Knockdown was confirmed by immunoblot analysis or qRT-PCR.

### Immunoblot analysis

For immunoblot analysis, 120,000 MZ-54 cells or LN-229 cells were seeded into 6-well plates. Cells were lysed with 2xSDS lysis buffer (137 mM Tris-HCl [Sigma-Aldrich, T1503] pH 6.8, 4% SDS [Serva Electrophoresis, 151-21-3], 20% glycerol [AppliChem 56-81-5], 1 mM protease inhibitor cocktail [Sigma-Aldrich, P8340]) and ultrasonic beats (3×10 times). The protein amount was determined with a Pierce BCA protein assay kit (Thermo Fisher Scientific, 23225). The SDS gels (8% - 15%) were loaded with 30 – 50 µg protein in 5X SDS loading buffer (250 mM Tris-HCl, pH 6.8, 10% SDS, 30% glycerol, 0.02% bromophenol blue [Sigma-Aldrich, B8026], 5% 2-mercaptoethanol [Sigma-Aldrich, M6250]) after heating the samples for 5 min at 95°C. Proteins were separated (85 V for 30 min, then 135 V) and blotted semi-dry (15 V, 35 min) onto a nitrocellulose membrane (Neolab, 260201396). The membranes were blocked with 5% milk (Carl Roth, T145.2) or 5% BSA (AppliChem, A1391.0500) in TBS (150 mM NaCl, 50 mM Tris, pH 7.5) + 0.05% Tween (AppliChem, A13890500) (TBS-T) for 1 h at room temperature, followed by incubation with the primary antibodies over night at 4°C: ATG5 (1:1000; Cell Signaling Technology, 2630s), ATG7 (1:500; Cell Signaling, 8558), RB1CC1 (1:1000; Proteintech, 17250-1-AP) MAP1LC3B/LC3B (1:1000; Thermo Fisher Scientific, PA1-16930), CTSB (1:1000; Cell Signaling, 31718), CTSD (1:1000; Cell Signaling, 2284), VCP (1:400; Abcam Biochemicals, ab11433), LAMP2 (1:1000; DSHB, H4B4), AKT1 (1:1000; Cell Signaling, 9272), phospho(Ser473)-AKT1 (1:2000; Cell Signaling, 4060), RPS6 (1:1000; Cell Signaling, 2317), phosphor(Ser240/244)-RPS6 (1:1000; Cell Signaling, 2215), TUBA4A (1:5000; Sigma-Aldrich, T6199) and GAPDH (1:10,000; Merck Millipore, CB1001). Secondary antibodies (Li-COR Biotech: goat anti-mouse, 926-32210 or 926-68070 and goat anti-rabbit, 926-32211 or 926-68071) were applied at a concentration of 1:10,000 for 1 h at room temperature. Detection and densiometric quantification of the secondary antibodies with the infrared dye was performed with the ‘Odyssey Infrared Imaging System’ (Li-COR Biosciences, Bad Homburg, Germany).

### Fluorescence microscopy

MZ-54 and LN-229 cells were seeded at 12,000 cells per chamberslide-well and fixed with 3.7% paraformaldehyde for 15 min, followed by three washing steps with PBS-Tween (0.01%; PBS-T). For cholesterol staining, cells were incubated with filipin III (1:100) in PBS for 2 h as described in the manufacturer’s protocol of the cholesterol assay kit (Abcam Biochemicals, ab133116). Filipin III was also added in all following incubation steps with the primary and secondary antibody to enable proper staining. For immunofluorescence staining, cells were permeabilized with 0.5% Triton in PBS for 5 min and blocked with 4 % BSA in PBS for 1 h at room temperature. Cells were incubated with an antibody against LAMP1 (anti-mouse; DSHB, H4A3s) over night at 4°C, followed by three washing steps with PBS-T and subsequent incubation with the secondary antibody (Alexa Fluor 488 goat anti-mouse IgG (H+L), Molecular Probes, A11029) for 1 h at room temperature. After another three washing steps, cover glasses were fixed with mounting medium with (Dianova, SCR-38448) or without DAPI (Thermo Fisher Scientific, TA-030-FM). Microscope images were taken with the Nikon Eclipse TE2000-S microscope (Nikon, Tokio, Japan) and NIS Elements AR 3.2 software at 60x magnification. Image analysis, optical zoom and overlays were performed with ImageJ Fiji (NIH, Bethesda, USA).

### Fractionation of cytosolic and lysosomal fraction for immunoblot analysis

Separation of the cytosolic fraction and the organelle fraction including fully intact lysosomes was performed by using digitonin as described by Adrian et al. [72]. MZ-54 control cells and MZ-54 *ATG5* KO1 were seeded at 2 - 3 *10^6^. Cell pellets were carefully resuspended with 500 µL chilled PBS, followed by addition of 500 µL cytosol extraction buffer (250 mM sucrose, 70 mM KCL, 137 mM NaCl, 4.3 mM Na_2_HPO_4_, 1.4 mM KH_2_PO_4_, pH 7.2. Freshly added: 100 µM PMSF, 10 µg/ml leupeptin, 2 µg/ml aprotinin, 50 µg/ml digitonin) and subsequent incubation for 15 min at room temperature on an overhead tumbler. Digitonin concentration and incubation time for the cytosol extraction was precisely titrated. For separation of the cytosol, cells were centrifuged at 1035 x g and 4°C for 5 min, supernatants were transferred into a new tube and centrifuged again for 30 sec at 1677 x g. Supernatants containing the cytosolic fraction were collected in a new tube. In the next step, the pellets from the first centrifugation step were resuspended in 300 µL lysis buffer (50 mM Tris/HCl pH 7.4, 150 mM NaCl, 2 mM EDTA, 2 mM EGTA, 0.2% Triton X-100, 0.3% NP-40; freshly added: 10 µg/ml leupeptin, 2 µg/ml aprotinin) and incubated for 5 min, followed by centrifugation for 10 min at 10,278 x g and 4°C. Collected supernatants were spinned down for 30 sec at 6708 x g and transferred into a new tube. The fractions contain various organelles including intact lysosomes. For immunoblot analysis of CTSB and CTSD release into the cytosol, 20 - 25 µg protein was used SDS-PAGE and Western blot.

### Cathepsin B assay

For fluorescence-based analysis of cathepsin B activity in the cytosolic fractions of MZ-54 cells, 30,000 cells were seeded into 24-well plates and treatments were performed on the next day. Cytosolic fractions were obtained by washing the cells with DPBS and subsequent incubation with 200 µl cytosol extract buffer containing a titrated amount of digitonin (250 mM sucrose, 20 mM HEPES, 10 mM KCL, 1.5 mM MgCl_2_, 1 mM EDTA, 1 mM EGTA; freshly added: Pefablock SC 0.5 mM, digitonin 22.5 µg/mL) for 15 min on ice while shaking slowly 73. Next, 100 µL cytosol extract was mixed with 75 µL L-cysteine buffer (352 mM KH_2_PO_4_, 48 mM Na_2_HPO_4_, 4 mM EDTA, pH 6; freshly added: 8 mM L-Cysteine) in a black 96-well plate and incubated at 37 °C for 10 min. After adding 75 µL cathepsin B reaction buffer (0.1% Brji 35 solution; freshly added: 0.02 mM Z-Arg-Arg-7-amido-4-methylcoumarin hydrochloride), the 96-well plate was immediately placed into a SPARK multimode microplate reader (Tecan, Männedorf, Switzerland) and fluorescence was measured every 5 min at 40°C over a time period of 2 h (Ex: 348 nm, Em: 440 nm).

### Flow cytometry analysis of cell death, autophagic flux and lipid-ROS

Cell death was measured by double staining with APC-ANXA5/annexin V (APC-annexin V; ex/em maxima 650/660; BD Pharmingen, 550475) and propidium iodide (PI; ex/em maxima ∼ 535/617 nm; Sigma-Aldrich, P4864) and subsequent flow cytometric analysis. Therefore, 30,000 cells were seeded into 24-well plates and treatments were performed on the next day. Subsequently, cells were pelletized and resuspended in 50 µl FACS buffer (10 mM HEPES, 140 mM NaCl, 5 mM CaCl_2,_ pH 7.4) with 0.8 µl APC-annexin V and 0.8 µg/mL PI. After 10 min incubation, measurements were performed with a FACS Accuri (BD Biosciences, Heidelberg, Germany) by using the FL2-A channel for the PI signal and the FL4-A channel for the APC-annexin V. DMSO-treated cells were gated APC-annexin V and PI negative. Overall cell death was calculated by summing up the percentage of only-APC-annexin V-positive, only-PI-positive and double-positive cells. The autophagic flux was measured by using the reporter construct pMRX-IP-GFP-LC3B-RFP-LC3BΔG [74] and subsequent flow cytometric analysis. For this, 30,000 cells stably transfected with the reporter construct were seeded into 24-well plates and treatments were performed on the next day. Subsequently, cells were harvested, pelletized and resuspended in 40-50 µL DPBS. After cleavage of the fluorescent probe by endogenous ATG4 proteases EGFP-LC3 is degraded in the lysosomes and mRFP1-LC3ΔG remains in the cytosol and serves as an internal control. Therefore, a decrease in the EGFP/mRFP1 ratio is indicative for an increase in autophagic flux. Measurements were performed by using the FL1-A channel for the EGFP signal and the FL3-A channel for the mRFP1 signal. For assessment of lipid-ROS levels, 28,000 cells were seeded into 24-well plates and stained with the lipid peroxidation sensor BODIPY™ 581/591 C11 (Thermo Fisher Scientific, D3861) at 5 µM for 30 min at 37°C. Subsequently, cells were harvested and resuspended in 50 µL DPBS. Cellular lipid-ROS levels were analyzed by quantification of the MFI signal of BODIPY™ 581/591 C11in the FL1-A channel.

### Assessment of gene expression by quantitative real-time polymerase chain reaction (qRT-PCR)

For determination of gene expression by qRT-PCR, 150,000 cells were seeded per 6-well or cells were directly collected from the cell culture flask. After washing the cell pellets with DPBS, RNA was isolated with the EXTRACTME total RNA Kit according to the manufacturer’s instructions (7Bioscience, EM09.1-250). Next, 1 – 2 µg RNA, superscript III reverse transcriptase (Thermo Fisher, 18080044) and random primers were used in a total volume of 20 µL for cDNA synthesis according to the manufacturer’s protocol. CDNA was diluted with 80-180 µL DEPC-H_2_O. For qRT-PCR, 5 µL cDNA, 1 μl 1xTaqMan Gene Expression Assay primer and 10 μl 1x FastStart Universal Probe Master-mix (Roche, 04913957001) were applied. Relative gene expression levels were calculated by using the comparative CT method [75]. Samples were normalized to the reference gene *TBP* (TATA-box binding protein). The following FAM (FAM-dye-labeled)-MGB (minor-grove-binding) primers were used: Hs00427620_m1 (*TBP*), Hs04195319_s1 (*CERS1*), Hs00371958_g1 (*CERS2*), Hs00908756_m1 (*CERS5*), Hs01372226_m1 (*CERS6*), Hs00186447_m1 (*DEGS1*), Hs01116902_m1 (*SPTLC1*), Hs00166163_m1 (*NOX2*), Hs00208,732_m1 (*ATG14*), (Thermo Fisher Scientific).

### Targeted analysis of sphingolipids and ceramides

For analysis of the sphingolipid composition, 3*10^6^ MZ-54 cells were seeded into T-175 flasks in octaplicates. Treatments were performed on the next day by using medium containing 2% FCS for loperamide treatments and 5% FCS for pimozide treatments and the respective controls to reduce background. Cells were harvested by using trypsin and washed four times with chilled DBPS, finally pelleted, resuspended in 20 µl, counted, weighed (i.e. difference versus empty tube) and snap frozen in liquid nitrogen and stored at -80°C until analysis.

Cell pellets were thawed and homogenized in 1 mL extraction buffer (citric acid 30 mM, disodium hydrogen phosphate 40 mM). Ten microliter of the homogenized sample were mixed with 200 µl extraction buffer and 20 µl of the internal standard solution containing deuterated sphingolipids and ceramides (all avanti polar lipids, Alabaster, AL, USA) as explained in detail in [76]. The mixture was extracted once with 600 µL methanol/chloroform/hydrochloric acid (15:83:2, v/v/v). The lower organic phase was evaporated at 45 °C under a gentle stream of nitrogen and reconstituted in 200 µL of tetrahydrofuran/water (9:1, v/v) with 0.2 formic acid and 10 mM ammonium formate.

For the preparation of calibration standards and quality control samples, 20 µl of a working solution were processed as explained above. Working solutions for the generation of calibrator and QC-samples were prepared as a mixture of all analytes by serial dilution using a mixture of tetrahydrofuran and chloroform. Quality control samples of three different concentration levels (low, middle, high) were run as initial and final samples of each run.

Amounts of sphingolipids were analyzed by liquid chromatography coupled to tandem mass spectrometry. An Agilent 1260 series binary pump (Agilent technologies, Waldbronn, Germany) equipped with a Zorbax Eclipse Plus C18 UHPLC column (50 mm x 2.1 mm ID, 1.8 μm, Agilent technologies, Waldbronn, Germany) was used for chromatographic separation. The column temperature was 55 °C. The HPLC mobile phases consisted of water with 0.2% formic acid and 10 mM ammonium formate (mobile phase A) and acetonitrile/isopropanol/acetone (50:30:20, v/v/v) with 0.2% formic acid (mobile phase B). For separation, a gradient program was used at a flow rate of 0.4 ml/min. The initial buffer composition 65% (A)/35% (B) was held for 0.6 min and then within 0.4 min, linearly changed to 35% (A)/65% (B) and held for 0.5 min. Within 3 min, (b) was further increased to 100% and held for 6.5 min. Subsequently, the composition was linearly changed within 0.5 min to 65% (A)/35% (B) and then kept for another 2.5 min. The total running time was 14 min, and the injection volume was 10 µl. To improve ionization, isopropyl alcohol was infused post-column using an isocratic pump at a flow rate of 0.1 ml/min. After every sample, sample solvent was injected for washing and re-equilibrating the analytical column using two short runs (4 min each).

The MS/MS analyses were performed using a triple quadrupole mass spectrometer QTrap 5500 (Sciex, Darmstadt, Germany) equipped with a Turbo V Ion Source operating in positive electrospray ionization mode as described in [76–78]. The analysis was done in Multiple Reaction Monitoring (MRM) mode. Precursor to product ion transitions (m/z) for analysis and internal standards are shown in the supplementary methods of [76]. Data Acquisition was done using Analyst Software V 1.6.2 and quantification was performed with MultiQuant Software 3.0.2 (both Sciex, Darmstadt, Germany), employing the internal standard method (isotope dilution mass spectrometry). Calibration curves were calculated by linear or quadratic regression with 1/x or 1/x2 weighting. Variations in accuracy of the calibration standards were less than 15% over the whole range of calibration, except for the lower limit of quantification, where a variation in accuracy of 20% was accepted. For the acceptance of the analytical run, the accuracy of the QC samples had to be between 85% and 115% of the nominal concentration for at least 67% of all QC samples.

### Proteomics

The proteome of MZ-54 cells was analyzed by using a Tandem Mass Tag (TMT) mass spectrometry. Therefore, 4*10^6^ cells were seeded per condition and treated on the next day in two independent experiments. Samples were washed two times with cold DPBS and harvested by using lysis buffer (2% SDS, 150 mM NaCl, 50 mM Tris pH 8.5, 40 mM chloroacetamide, 5 mM TCEP; freshly added: 1 tablet Phosphostop [Roche, 4906845001] and 1 tablet cOmplete Protease inhibitor cocktail [Roche, 4693159001]/10 mL lysis buffer) and a cell scraper. Next, samples were boiled for 10 min at 95°C, sonicated for 2 min (1 sec on, 1 sec off, 45% amplitude) and boiled again for 5 min at 95°C. For methanol/chloroform precipitation, samples were mixed with four parts ice-cold methanol and one part ice-cold chloroform, followed by addition of three parts of ice-cold, deionized H_2_O and subsequent centrifugation at 4000 x g and 4 °C for 25 min. After removing the top-layer and adding three parts of ice-cold methanol, samples were again vortexed and centrifuged for 10 min at 4000 x g and 4°C. In the next step, the remaining protein pellet was washed with three parts ice-cold methanol and spinned down for 10 min at 4°C and 4000 x g. After another washing step with 1 mL ice-cold methanol, samples were centrifuged for 5 min at 15,000 – 20,000 g and 4°C and pellets were dried. The protein pellets were dissolved in 1 mL digestion buffer (8 M urea, 50 mM Tris pH 8.0) by heating to 37°C for 30 min and subsequent sonication. The protein amount was quantified with the Roti Quant colorimetric protein assay according to the manufacturer’s instructions (Carl Roth, Ko15.1). For digestion, 100 µg proteins were diluted 1:1 with 50 mM Tris pH 8.5 and 1 µg endoproteinase Lys-C (Wako Chemicals, 125-02543), followed by incubation for 3 h at 37°C while shaking. Next, protein samples were diluted 1:1 in EPPS buffer (10 mM EPPS [Sigma-Aldrich, E9502], 1 mM CaCl_2_), and digested with 1 µg trypsin (Promega, V5113) over night at 37°C while shaking. Digestion was stopped with 1% trifluoroacetic acid (TFA) (Sigma-Aldrich, 302031). Desalting of the peptides was carried out by using reverse-phase Sep-Pak tC18 cartridges with 50 mg sorbent (Waters Corporation, WAT054960) connected to a vacuum manifold. After washing and equilibration, the peptides were loaded, followed by a washing step with 3 mL 1% acetonitrile (ACN; VWR, 83640-290)/0.1% TFA and elution with 500 µL 40% ACN and 500 µL 60% ACN. Peptides were concentrated for 2 h in a vacuum concentrator (Eppendorf Vacuum Concentrator plus) at 30°C and diluted to 100 µL with H_2_O. For peptide quantification, a Micro BCA assay was performed according to the manufacturer’s instructions (Thermo Fisher Scientific, 23235). For TMT labeling of the peptides, an amount of 10 µg peptides was dried and resuspended in 10 µL 0.2 M EPPS (pH 8.2) / 20% ACN and the respective TMT10 reagent (Thermo Fisher Scientific, 1862804) in a TMT:peptide ratio of 2.5:1 was added. After 1 h incubation, the labeling reaction was quenched with hydroxylamine (Sigma-Aldrich, 438227) at a final concentration of 0.5% and incubated for 15 min before acidification with TFA (1% final concentration). After verification of labelling efficiency (>99%) and equal mixing by liquid chromatography – mass spectrometry (LC-MS), all samples were pooled, dried at 30°C in a vacuum concentrator and fractionated using a Pierce High pH Reverse-Phase Peptide Fractionation Kit (Thermo Fisher Scientific, 84868) according to the manufacturer’s instructions, except that 16 fractions were generated using 7.5, 10, 12.5, 15, 17.5, 20, 22.5, 25, 27.5, 30, 32.5, 35, 37.5, 42.5, 47.5 and 60% ACN, which were then pooled to yield 8 final fractions (F1+F9, F2+F10, …, F8+F16). Peptide fractions were evaporated to dryness and finally, samples were stored at -80°C until mass spectrometry analysis, for which they were resuspended in LC-MS grade water containing 2% ACN and 0.1% formic acid. Peptides were separated on an easy nLC 1200 (ThermoFisher) through a 75µm ID fused-silica column, which was packed in-house with 1.9 µm C18 particles (ReproSil-Pur, Dr. Maisch) to an approximate length of 20 cm and kept at 45 °C using an integrated column oven (Sonation). Peptides were eluted by a linear gradient from 5-38% acetonitrile over 150 minutes and directly sprayed into a QExactive HF mass-spectrometer equipped with a nanoFlex ion source (ThermoFisher Scientific) at a spray voltage of 2.3 kV. Full scan MS spectra (350-1400 m/z) were acquired at a resolution of 120,000 at m/z 200, a maximum injection time of 100 ms and an AGC target value of 3 x 10^6^ charges. Up to 12 most intense peptides per full scan were isolated using a 0.8 Th window and fragmented using higher energy collisional dissociation (HCD) applying a normalized collision energy of 35. MS/MS spectra were acquired with a resolution of 45,000 at m/z 200, a maximum injection time of 128 ms and an AGC target value of 1 x 10^5^. Ions with charge states of 1 and > 6 as well as ions with unassigned charge states were not considered for fragmentation. Dynamic exclusion was set to 20 s to minimize repeated sequencing of already acquired precursors.

### Analysis of mass spectrometry data

Acquired raw files were analyzed with MaxQuant (version 1.6.1.0). Spectra were matched against a database containing all human entries including isoforms contained in SwissProt (taxonomy ID 9606, 42144 sequences, downloaded 2017-09-02). Mostly, default parameters for a “reporter ion MS2”-type of experiment were used, except: protein groups required at least 1 unique peptide to be reported, second peptide search was disabled and only spectra with a PIF >0.7 as well as also unmodified counterpart peptides were used for quantification. Reporter ion intensities were corrected for isotopic impurities as specified by the manufacturer and median-normalized in R using the Normalyzer package 79,[80]. Gene ontology (GO) term analysis was performed with Perseus [81] and String (string-db.org; PMID: 25352553; version 11.0). The mass spectrometry proteomics data have been deposited to the ProteomeXchange Consortium (http://proteomecentral.proteomexchange.org) via the PRIDE partner repository [82] with the dataset identifier PXD015074.

Reviewer account details:

Username

Password: agYAfU7G

### Statistics

Statistical analysis of proteome data was performed in Perseus. Only proteins with P ≤ 0.01 based on a Student’s t-test were considered significant. GraphPad Prism 7 (GraphPad Software, La Jolla CA, USA) was used for statistical analysis of all other data. To determine if the data set is well-modeled by a normal distribution, a Shapiro Wilk normality test was performed. In case of normally distributed data, the data set was analyzed with a one- or two-way ANOVA. For non-normally distributed data, the non-parametric Mann-Whitney U-test or Kruskal-Wallis test was used. The minimum level of statistical significance was set at P ≤ 0.05 and significances are depicted as P ≤ 0.05: *, P ≤ 0.01: **, P ≤ 0.001: *** or n.s.: not significant between control and treated cells or as indicated with brackets.

---

**Figure S1:**
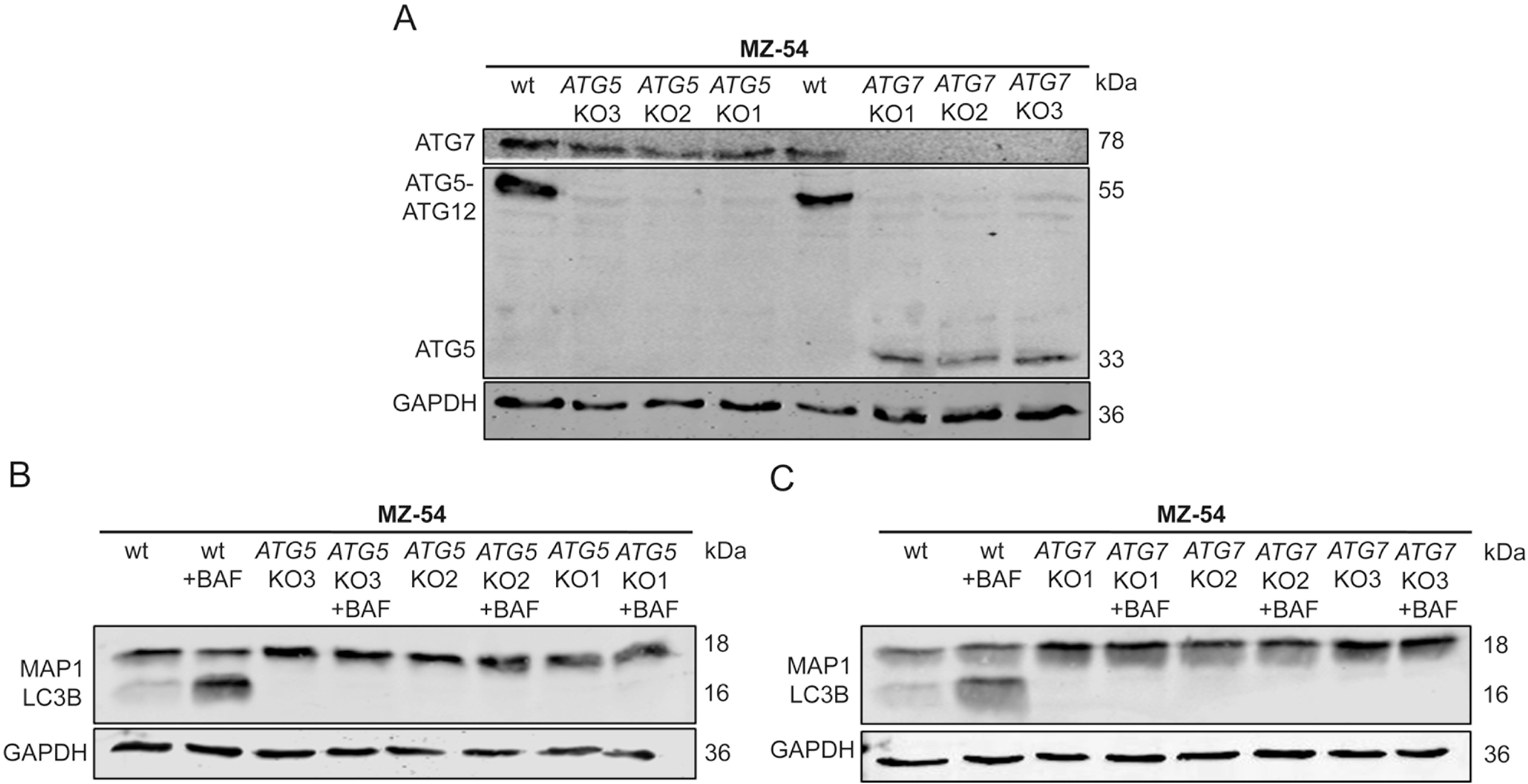
Determination of autophagy inhibition by *ATG5* and *ATG7* knockout. (**A-C**) Immunoblot analysis of MAP1LC3B-I and -II (**B, C**), ATG5 and ATG7 expression (**A**). GAPDH was used as housekeeper. (**B, C**) MZ-54 wt cells and the respective *ATG5* and *ATG7* knockouts were treated with 10 nM bafilomycin A_1_ (BAF) for 4 h.

**Figure S2:**
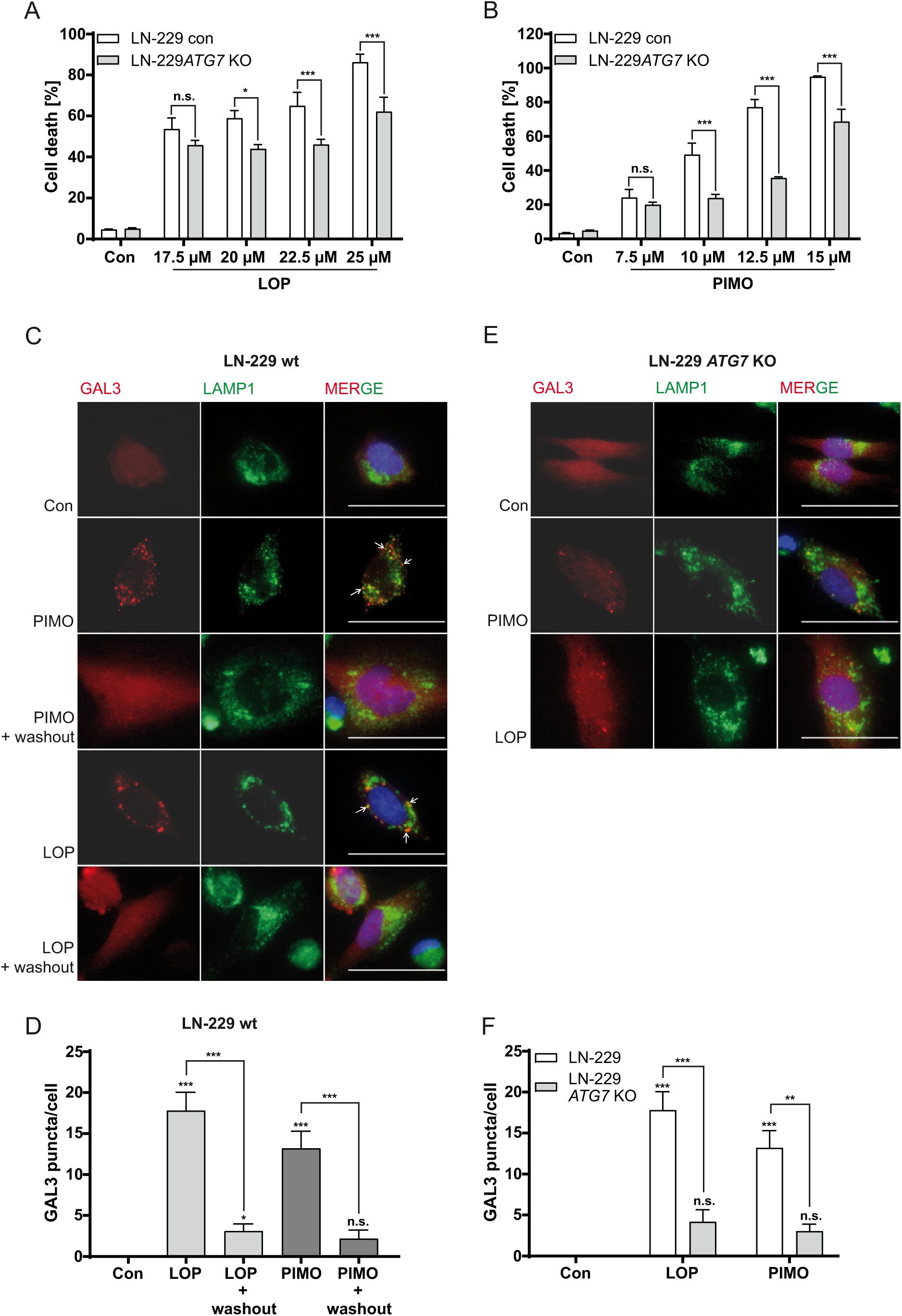
Determination of autophagic cell death and lysosomal membrane permeabilization after pimozide and loperamide treatment in LN-229 cells. (**A, B**) Flow cytometric analysis of cell death by measurement of only-APC-annexin V-positive, only-PI-positive and double-positive cells. LN-229 wt cells and CRISPR/Cas9 *ATG7* KO cells were treated with different concentrations of loperamide (LOP) (**A**), pimozide (PIMO) (**B**) or DMSO (Con) for 48 h. Data show mean + SEM of at least three experiments with three replicates and 5,000 – 10,000 cells measured in each sample. Statistical significances were calculated with a two-way ANOVA. (**C-F**) Microscopic assessment of lysosmal membrane permeabilization by evaluating the mCherry-GAL3 puncta formation and colocalization (depicted with arrows) with the lysosomal marker LAMP1. LN-229 wt (**C**) and *ATG7* KO cells (**E**) stably transfected with pmCherry-GAL3 were treated with 12.5 µM pimozide (PIMO), 22.5 µM loperamide (LOP) or DMSO (Con) for 16 h. For washout experiments shown in **C**, the drug-containing media were removed after 16 h and fresh medium was added for additional 24 h. Three independent experiments were performed and three to eight images were taken at 60x magnification in each experiment (scale bar=50 µM). (**D, F**) Quantification of mCherry-GAL3 puncta/cell. In total 24 – 51 cells were analyzed per condition. Data represent mean + SEM of three independent experiments and three to eight images per experiment. Statistical significances were analyzed with a Kruskal-Wallis test.

**Figure S3.**
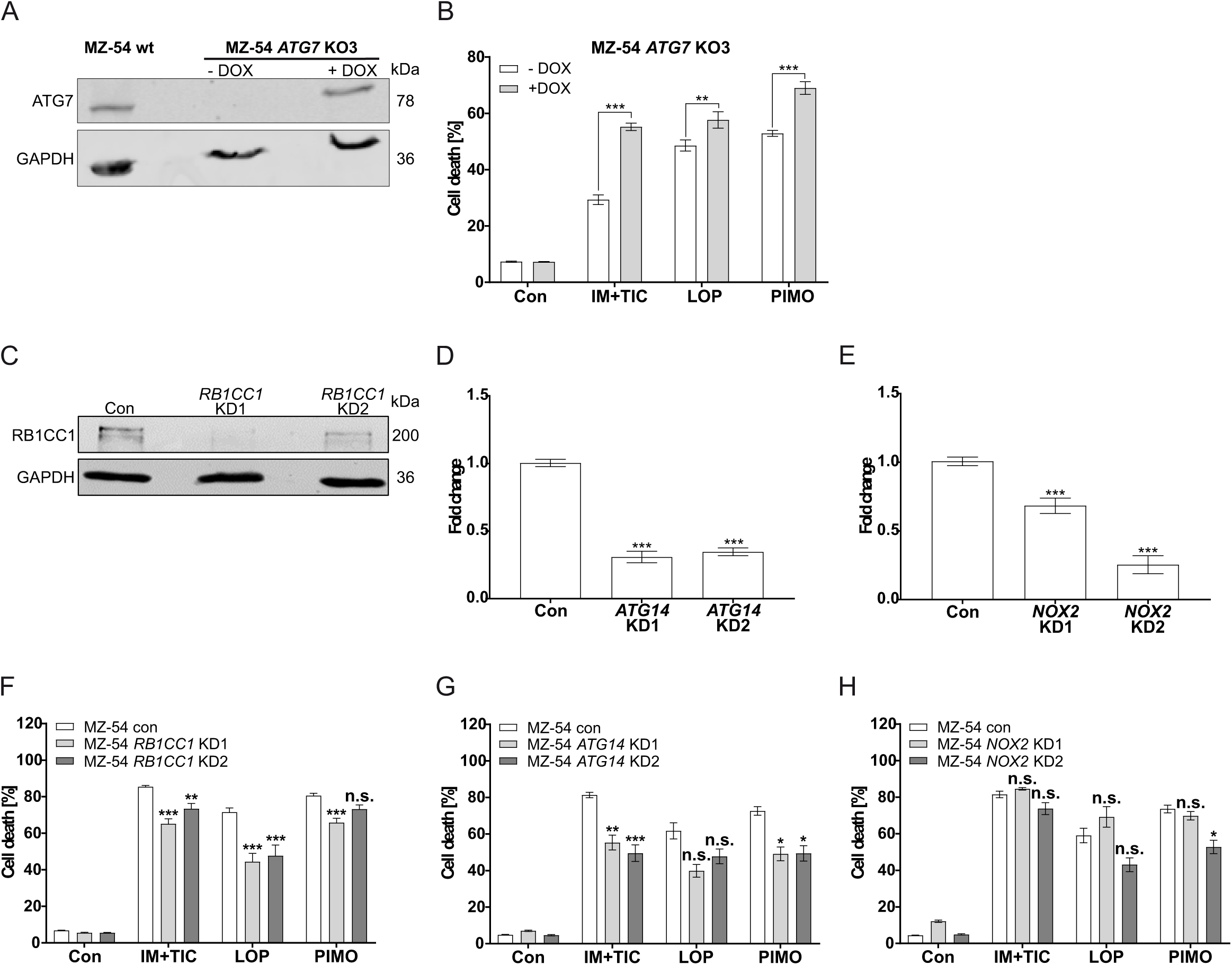
Re-expression of *ATG7* in MZ-54 *ATG7* knockout cells and assessment of LAP and autophagy involvement in imipramine + ticlopidine-, pimozide- and loperamide-induced cell death. (A) Validation of the re-expression of *ATG7* in *ATG7* KO cells transduced with the TRE3G-*ATG7* and Tet3G plasmids by western blot. Cells were treated with 1 µg/mL doxycycline (DOX) for 24 h in order to induce the expression of *ATG7*. The experiment was repeated two times. **(B)** Determination of cell death by flow cytometric quantification of only-APC-annexin V-positive, only-PI-positive and double-positive cells. MZ-54 *ATG7* KO cells re-expressing *ATG7* after the addition of 1 µg/mL doxycycline (DOX) were treated with 20 µM imipramine + 100 µM ticlopidine (IM+TIC), 12.5 µM loperamide (LOP), 12.5 µM pimozide (PIMO) or DMSO (Con) for 48 h. Data show mean + SEM of four independent experiments with three replicates and 5,000 – 10,000 cells measured in each sample. Statistical significances were analyzed with a two-way ANOVA. (**C-E**) Knockdown validation of the autophagy specific genes *RB1CC1* and *ATG14* and the LAP specific gene *NOX2* in MZ-54 cells. Knockdown was approved either by western blot (**C**) or by qRT-PCR (**D, E**). (**C**) Experiments were repeated two times. (**D, E**) Data represent mean + SEM of two - five independent experiments with three replicates. Statistical significances were calculated with a Kruskal-Wallis test. (**F-H**) Assessment of cell death as described above. MZ-54 control (con) and *RB1CC1*, *ATG14* and *NOX2* KD cells were treated with 20 µM imipramine + 100 µM ticlopidine (IM+TIC), 12.5 µM loperamide (LOP), 12.5 µM pimozide (PIMO) or DMSO (Con) for 48 h. Data show mean + SEM of at least four independent experiments with three replicates and 5,000 – 10,000 cells measured in each sample. Statistical significances were analyzed with a two-way ANOVA (**F**) or a Kruskal-Wallis test (**G, H**).

**Figure S4.**
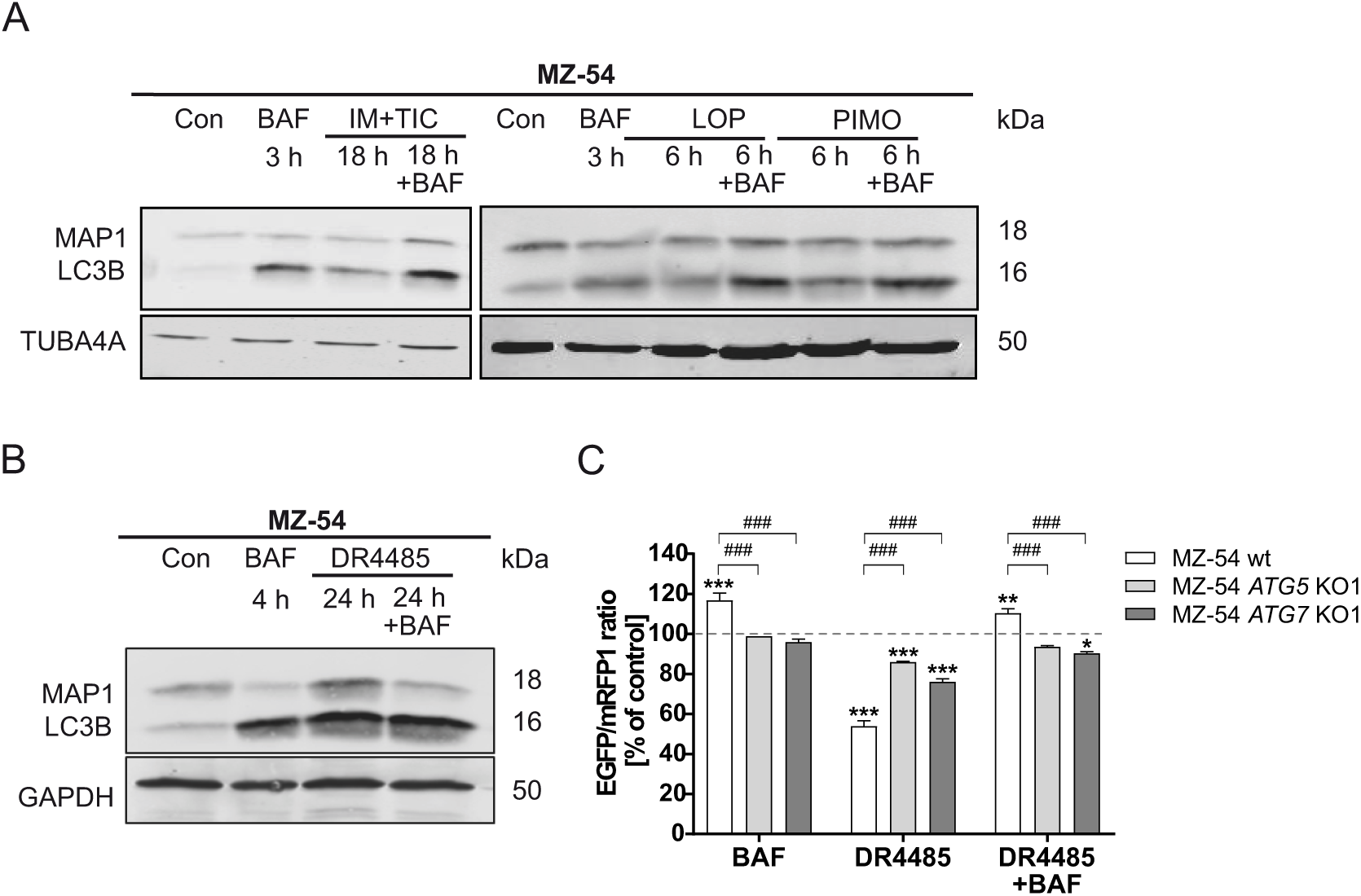
Determination of the autophagic flux after imipramine + ticlopidine, loperamide, pimozide and DR4485 treatment. (**A, B**) Immunoblot analysis of MAP1LC3B-I and MAP1LC3B-II expression. TUBA4A (**A**) or GAPDH (**B**) were used as housekeepers. (**A**) MZ-54 cells were exposed to 20 µM imipramine + 100 µM ticlopidine (IM+TIC, 18 h), 7.5 µM loperamide (LOP, 6 h), 7.5µM pimozide (PIMO, 6 h) or DMSO (Con, 18 h) alone or in combination with bafilomycin A1 (BAF). Bafilomycin A1 was added to the cells 3 h before harvest. (**B**) MZ-54 cells were treated with 6 µM DR4485 for 24 h in the presence or absence of bafilomycin A1 (BAF) for the last 4 h. Experiments were repeated two times. (**C**) Flow cytometric analysis of the autophagic flux by measurement of the EGFP and mRFP1 signal. MZ-54 wt cells and *ATG5* and *ATG7* KO cells were exposed to 4 µM DR4485 alone or in combination with bafilomycin A1 (BAF) for 24 h. Data show mean + SEM of three independent experiments with three replicates and 5,000 – 10,000 cells measured in each sample. Statistical significances were calculated with a two-way ANOVA. Significances between wt and KO cells are depicted as P ≤ 0.05: #, P ≤ 0.01: ## or P ≤ 0.001: ###.

**Figure S5.**
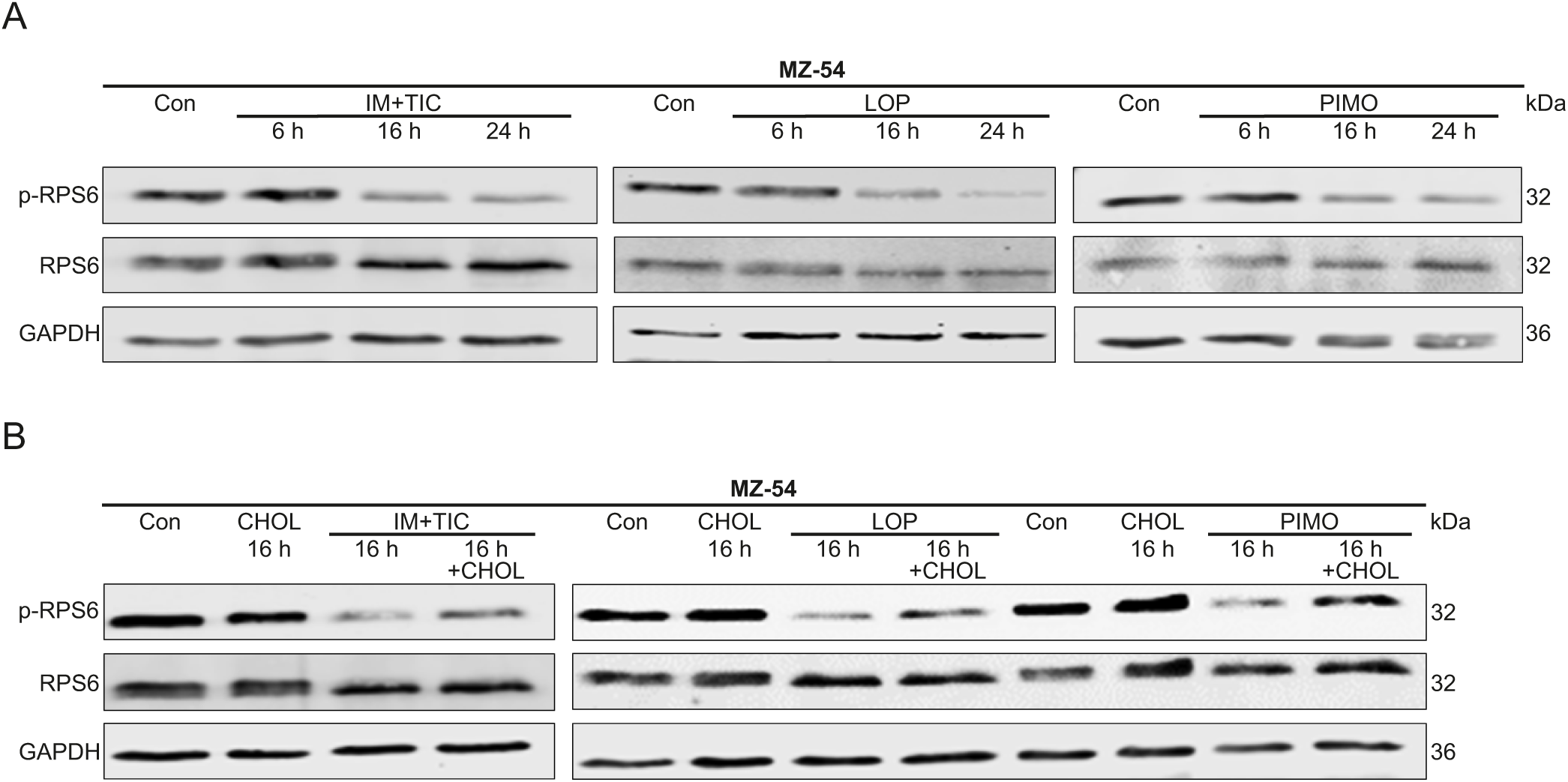
Analysis of ribosomal protein S6 kinase activity in imipramine + ticlopidine, loperamide and pimozide treated MZ-54 cells. (**A, B**) Immunoblot analysis of phosphorylated ribosomal protein S6 (RPS6) and RPS6. GAPDH was used as housekeeper. MZ-54 cells were exposed to 20 µM imipramine + 100 µM ticlopidine (IM+TIC), 15 µM loperamide (LOP), 15 µM pimozide (PIMO) with or without 10 µg/mL cholesterol–methyl-β-cyclodextrin for the indicated time periods (6 h, 16 h, 24 h). DMSO was used as control (Con, 24 h (**A**), 16 h (**B**)). Experiments were repeated at least two times.

**Figure S6.**
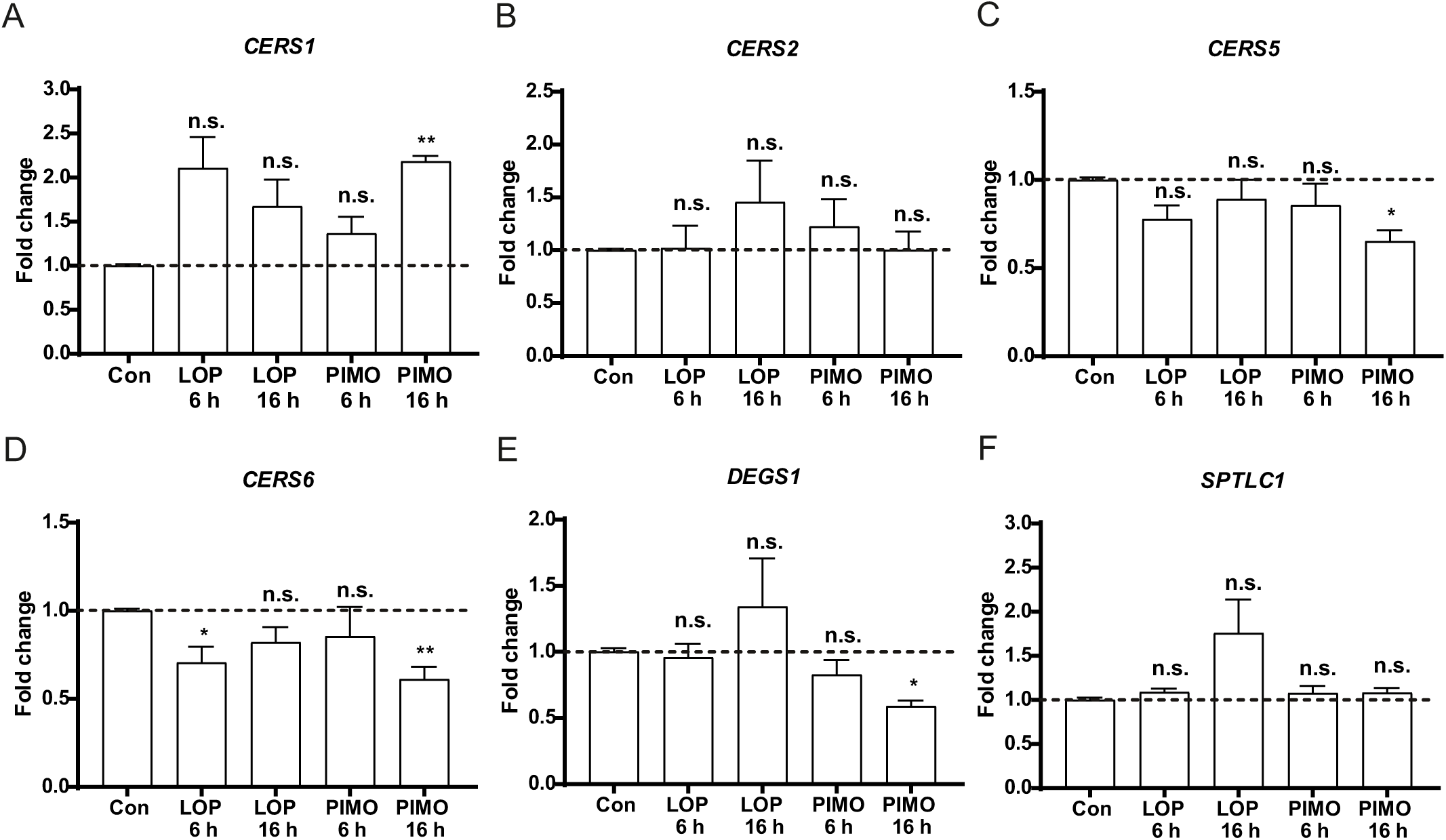
Expression analysis of enzymes involved in *de novo* ceramide synthesis. (**A-F**) Expression levels of *CERS1, CERS2, CERS5, CERS6, DEGS1* and S*PTLC*1 mRNA normalized to *TBP* were analyzed by qRT-PCR. MZ-54 cells were treated with 12.5 µM loperamide (LOP), 12.5 µM pimozide (PIMO) or DMSO (Con) for 6 h and 16 h. Data represent mean + SEM of three independent experiments with three technical replicates. Statistical analysis was done with a Kruskal-Wallis test.

## Acknowledgements

This study was supported by the German Research Foundation (Deutsche Forschungsgemeinschaft, SFB 1177 on selective autophagy to D.K., S.F and C.M; SFB1039, subproject A3 to I.T.)

## Notes

http://proteomecentral.proteomexchange.org

### Competing Interest Statement

The authors have declared no competing interest.

## References

[1] Mariño G, Niso-Santano M, Baehrecke EH, et al. Self-consumption. Nat Rev Mol Cell Biol. 2014; 15(2):81–94.

[2] Klionsky DJ, Cuervo AM, Dunn WA, et al. How shall I eat thee? Autophagy. 2007; 3(5):413–6.

[3] Liu Y-L, Yang P-M, Shun C-T, et al. Autophagy potentiates the anti-cancer effects of the histone deacetylase inhibitors in hepatocellular carcinoma. Autophagy. 2010; 6(8):1057– 65.

[4] Dasari SK, Bialik S, Levin-Zaidman S, et al. Signalome-wide RNAi screen identifies GBA1 as a positive mediator of autophagic cell death. Cell Death Differ. 2017; 24(7):1288–302.

[5] Segala G, David M, Medina P de, et al. Dendrogenin A drives LXR to trigger lethal autophagy in cancers. Nat Commun. 2017; 8(1):1903.

[6] Fulda S, Kögel D. Cell death by autophagy. Oncogene. 2015; 34(40):5105–13.

[7] Thakkar JP, Dolecek TA, Horbinski C, et al. Epidemiologic and molecular prognostic review of glioblastoma. Cancer Epidemiol Biomark Prev. 2014; 23(10):1985–96.

[8] Davis ME. Glioblastoma. Clin J Oncol Nurs. 2016; 205 Suppl:S2-8.

[9] Wagenknecht B, Glaser T, Naumann U, et al. Expression and biological activity of X-linked inhibitor of apoptosis (XIAP) in human malignant glioma. Cell Death Differ. 1999; 6(4):370–6.

[10] Jiang Z, Zheng X, Rich KM. Down-regulation of Bcl-2 and Bcl-xL expression with bispecific antisense treatment in glioblastoma cell lines induce cell death. J Neurochem. 2003; 84(2):273–81.

[11] Voss V, Senft C, Lang V, et al. The pan-Bcl-2 inhibitor (-)-gossypol triggers autophagic cell death in malignant glioma. Mol Cancer Res. 2010; 8(7):1002–16.

[12] Salazar M, Carracedo A, Salanueva ÍJ, et al. Cannabinoid action induces autophagy-mediated cell death through stimulation of ER stress in human glioma cells. J Clin Investig. 2009; 119(5):1359–72.

[13] Shchors K, Massaras A, Hanahan D. Dual Targeting of the Autophagic Regulatory Circuitry in Gliomas with Repurposed Drugs Elicits Cell-Lethal Autophagy and Therapeutic Benefit. Cancer Cell. 2015; 28(4):456–71.

[14] Kögel D, Fulda S, Mittelbronn M. Therapeutic Exploitation of Apoptosis and Autophagy for Glioblastoma. Anti-Cancer Agents Med Chem. 2010; 10(6):438–49.

[15] Linder B, Kögel D. Autophagy in Cancer Cell Death. Biology (Basel). 2019; 8(4).

[16] Galluzzi L, Vitale I, Aaronson SA, et al. Molecular mechanisms of cell death. Cell Death Differ. 2018; 25(3):486–541.

[17] Bialik S, Dasari SK, Kimchi A. Autophagy-dependent cell death - where, how and why a cell eats itself to death. J Cell Sci. 2018; 131(18).

[18] Das G, Shravage BV, Baehrecke EH. Regulation and function of autophagy during cell survival and cell death. Cold Spring Harb Perspect Biol. 2012; 4(6).

[19] Zielke S, Meyer N, Mari M, et al. Loperamide, pimozide, and STF-62247 trigger autophagy-dependent cell death in glioblastoma cells. Cell Death Dis. 2018; 9(10):994.

[20] Church J, Fletcher EJ, Abdel-Hamid K, et al. Loperamide blocks high-voltage-activated calcium channels and N-methyl-D-aspartate-evoked responses in rat and mouse cultured hippocampal pyramidal neurons. Mol Pharmacol. 1994; 45(4):747–57.

[21] DeHaven-Hudkins DL, Burgos LC, Cassel JA, et al. Loperamide (ADL 2-1294), an opioid antihyperalgesic agent with peripheral selectivity. J Pharmacol Exp Ther. 1999; 289(1):494–502.

[22] Freedman SB, Patel S, Marwood R, et al. Expression and pharmacological characterization of the human D3 dopamine receptor. J Pharmacol Exp Ther. 1994; 268(1):417–26.

[23] Elmaci I, Altinoz MA. Targeting the cellular schizophrenia. Likely employment of the antipsychotic agent pimozide in treatment of refractory cancers and glioblastoma. Crit Rev Oncol Hematol. 2018; 128:96–109.

[24] Heckmann BL, Green DR. LC3-associated phagocytosis at a glance. J Cell Sci. 2019; 132(5).

[25] Kast RE. Glioblastoma chemotherapy adjunct via potent serotonin receptor-7 inhibition using currently marketed high-affinity antipsychotic medicines. Br J Pharmacol. 2010; 161(3):481–7.

[26] McMaster CR. Lipid metabolism and vesicle trafficking. Biochem Cell Biol. 2001; 79(6):681–92.

[27] Amaya C, Fader CM, Colombo MI. Autophagy and proteins involved in vesicular trafficking. FEBS Lett. 2015; 589(22):3343–53.

[28] Noda T. Autophagy in the context of the cellular membrane-trafficking system. Biochem Soc Trans. 2017; 45(6):1323–31.

[29] Szklarczyk D, Franceschini A, Wyder S, et al. STRING v10: protein–protein interaction networks, integrated over the tree of life. Nucleic Acids Res. 2014; 43 Database issue:D447–52.

[30] Kristiana I, Sharpe LJ, Catts VS, et al. Antipsychotic drugs upregulate lipogenic gene expression by disrupting intracellular trafficking of lipoprotein-derived cholesterol. Pharmacogenomics J. 2010; 10(5):396–407.

[31] Horton JD, Goldstein JL, Brown MS. SREBPs. J Clin Investig. 2002; 109(9):1125–31.

[32] Schulze H, Sandhoff K. Lysosomal lipid storage diseases. Cold Spring Harb Perspect Biol. 2011; 3(6).

[33] Santos-Lozano A, Villamandos García D, Sanchis-Gomar F, et al. Niemann-Pick disease treatment. Ann Transl Med. 2015; 3(22):360.

[34] Kristiana I, Yang H, Brown AJ. Different kinetics of cholesterol delivery to components of the cholesterol homeostatic machinery. Biochim Biophys Acta. 2008; 178111–12:724–30.

[35] Lieberman AP, Puertollano R, Raben N, et al. Autophagy in lysosomal storage disorders. Autophagy. 2012; 8(5):719–30.

[36] Xu J, Dang Y, Ren YR, et al. Cholesterol trafficking is required for mTOR activation in endothelial cells. Proc Natl Acad Sci U S A. 2010; 107(10):4764–9.

[37] Kornhuber J, Muehlbacher M, Trapp S, et al. Identification of novel functional inhibitors of acid sphingomyelinase. PLoS One. 2011; 6(8):e23852.

[38] Kornhuber J, Tripal P, Reichel M, et al. Identification of new functional inhibitors of acid sphingomyelinase using a structure-property-activity relation model. J Med Chem. 2008; 51(2):219–37.

[39] Kölzer M, Werth N, Sandhoff K. Interactions of acid sphingomyelinase and lipid bilayers in the presence of the tricyclic antidepressant desipramine. FEBS Lett. 2004; 5591–3:96– 8.

[40] Gabandé-Rodríguez E, Boya P, Labrador V, et al. High sphingomyelin levels induce lysosomal damage and autophagy dysfunction in Niemann Pick disease type A. Cell Death Differ. 2014; 21(6):864–75.

[41] Klutzny S, Lesche R, Keck M, et al. Functional inhibition of acid sphingomyelinase by Fluphenazine triggers hypoxia-specific tumor cell death. Cell Death Dis. 2017; 8(3):e2709.

[42] Meyer N, Zielke S, Michaelis JB, et al. AT 101 induces early mitochondrial dysfunction and HMOX1 (heme oxygenase 1) to trigger mitophagic cell death in glioma cells. Autophagy. 2018; 14(10):1693–709.

[43] Wang Z, Wen L, Zhu F, et al. Overexpression of ceramide synthase 1 increases C18-ceramide and leads to lethal autophagy in human glioma. Oncotarget. 2017; 8(61):104022–36.

[44] Scarlatti F, Bauvy C, Ventruti A, et al. Ceramide-mediated macroautophagy involves inhibition of protein kinase B and up-regulation of beclin 1. J Biol Chem. 2004; 279(18):18384–91.

[45] Ordoñez R, Fernández A, Prieto-Domínguez N, et al. Ceramide metabolism regulates autophagy and apoptotic cell death induced by melatonin in liver cancer cells. J Pineal Res. 2015; 59(2):178–89.

[46] Hernández-Tiedra S, Fabriàs G, Dávila D, et al. Dihydroceramide accumulation mediates cytotoxic autophagy of cancer cells via autolysosome destabilization. Autophagy. 2016; 12(11):2213–29.

[47] Serrano-Puebla A, Boya P. Lysosomal membrane permeabilization in cell death. Ann N Y Acad Sci. 2016; 1371(1):30–44.

[48] Papadopoulos C, Kirchner P, Bug M, et al. VCP/p97 cooperates with YOD1, UBXD1 and PLAA to drive clearance of ruptured lysosomes by autophagy. EMBO J. 2017; 36(2):135–50.

[49] Chauhan S, Kumar S, Jain A, et al. TRIMs and Galectins Globally Cooperate and TRIM16 and Galectin-3 Co-direct Autophagy in Endomembrane Damage Homeostasis. Dev Cell. 2016; 39(1):13–27.

[50] Maejima I, Takahashi A, Omori H, et al. Autophagy sequesters damaged lysosomes to control lysosomal biogenesis and kidney injury. EMBO J. 2013; 32(17):2336–47.

[51] Wiklund ED, Catts VS, Catts SV, et al. Cytotoxic effects of antipsychotic drugs implicate cholesterol homeostasis as a novel chemotherapeutic target. Int J Cancer. 2010; 126(1):28–40.

[52] Kuzu OF, Gowda R, Noory MA, et al. Modulating cancer cell survival by targeting intracellular cholesterol transport. Br J Cancer. 2017; 117(4):513–24.

[53] Villa GR, Hulce JJ, Zanca C, et al. An LXR-Cholesterol Axis Creates a Metabolic Co-Dependency for Brain Cancers. Cancer Cell. 2016; 30(5):683–93.

[54] Petersen NHT, Olsen OD, Groth-Pedersen L, et al. Transformation-associated changes in sphingolipid metabolism sensitize cells to lysosomal cell death induced by inhibitors of acid sphingomyelinase. Cancer Cell. 2013; 24(3):379–93.

[55] Boya P, Kroemer G. Lysosomal membrane permeabilization in cell death. Oncogene. 2008; 27(50):6434–51.

[56] Yamane M, Moriya S, Kokuba H. Visualization of ceramide channels in lysosomes following endogenous palmitoyl-ceramide accumulation as an initial step in the induction of necrosis. Biochem Biophys Rep. 2017; 11:174–81.

[57] Gulbins A, Schumacher F, Becker KA, et al. Antidepressants act by inducing autophagy controlled by sphingomyelin-ceramide. Mol Psychiatry. 2018.

[58] Yang W-E, Ho C-C, Yang S-F, et al. Cathepsin B Expression and the Correlation with Clinical Aspects of Oral Squamous Cell Carcinoma. PLoS One. 2016; 11(3):e0152165.

[59] Perera RM, Stoykova S, Nicolay BN, et al. Transcriptional control of autophagy-lysosome function drives pancreatic cancer metabolism. Nature. 2015; 524(7565):361–5.

[60] Sun T, Jiang D, Zhang L, et al. Expression profile of cathepsins indicates the potential of cathepsins B and D as prognostic factors in breast cancer patients. Oncol Lett. 2016; 11(1):575–83.

[61] Gondi CS, Rao JS. Cathepsin B as a cancer target. Expert Opin Ther Targets. 2013; 17(3):281–91.

[62] Fehrenbacher N, Bastholm L, Kirkegaard-Sørensen T, et al. Sensitization to the lysosomal cell death pathway by oncogene-induced down-regulation of lysosome-associated membrane proteins 1 and 2. Cancer Res. 2008; 68(16):6623–33.

[63] Domagala A, Fidyt K, Bobrowicz M, et al. Typical and Atypical Inducers of Lysosomal Cell Death. Int J Mol Sci. 2018; 19(8).

[64] Ulbrich K, Knobloch T, Kreuter J. Targeting the insulin receptor. J Drug Target. 2011; 19(2):125–32.

[65] Faraz S, Pannullo S, Rosenblum M, et al. Long-term survival in a patient with glioblastoma on antipsychotic therapy for schizophrenia. Ther Adv Med Oncol. 2016; 8(6):421–8.

[66] Lee J-K, Nam D-H, Lee J. Repurposing antipsychotics as glioblastoma therapeutics. Oncol Lett. 2016; 11(2):1281–6.

[67] Tan SK, Jermakowicz A, Mookhtiar AK, et al. Drug Repositioning in Glioblastoma. Front Pharmacol. 2018; 9:218.

[68] Sánchez-Wandelmer J, Dávalos A, La Peña G de, et al. Haloperidol disrupts lipid rafts and impairs insulin signaling in SH-SY5Y cells. Neuroscience. 2010; 167(1):143–53.

[69] Canfrán-Duque A, Casado ME, Pastor O, et al. Atypical antipsychotics alter cholesterol and fatty acid metabolism in vitro. J Lipid Res. 2013; 54(2):310–24.

[70] Hetschko H, Voss V, Seifert V, et al. Upregulation of DR5 by proteasome inhibitors potently sensitizes glioma cells to TRAIL-induced apoptosis. FEBS J. 2008; 275(8):1925– 36.

[71] Ran FA, Hsu PD, Wright J, et al. Genome engineering using the CRISPR-Cas9 system. Nat Protoc. 2013; 8(11):2281–308.

[72] Adrain C, Creagh EM, Martin1 SJ. Apoptosis-associated release of Smac/DIABLO from mitochondria requires active caspases and is blocked by Bcl-2. EMBO J. 2001; 20(23):6627–36.

[73] Platt F, Platt N, eds. Lysosomes and lysosomal diseases. London, San Diego, Waltham, Oxford: Elsevier/Academic Press, 2015.

[74] Kaizuka T, Morishita H, Hama Y, et al. An Autophagic Flux Probe that Releases an Internal Control. Mol Cell. 2016; 64(4):835–49.

[75] Schmittgen TD, Livak KJ. Analyzing real-time PCR data by the comparative C(T) method. Nat Protoc. 2008; 3(6):1101–8.

[76] Brunkhorst-Kanaan N, Klatt-Schreiner K, Hackel J, et al. Targeted lipidomics reveal derangement of ceramides in major depression and bipolar disorder. Metab Clin Exp. 2019; 95:65–76.

[77] Zschiebsch K, Fischer C, Pickert G, et al. Tetrahydrobiopterin Attenuates DSS-evoked Colitis in Mice by Rebalancing Redox and Lipid Signalling. J Crohns Colitis. 2016; 10(8):965–78.

[78] Lötsch J, Schiffmann S, Schmitz K, et al. Machine-learning based lipid mediator serum concentration patterns allow identification of multiple sclerosis patients with high accuracy. Sci Rep. 2018; 8(1):14884.

[79] R Foundation for Statistical Computing. R: A Language and Environment for Statistical Computing: R Core Team, 2014.

[80] Chawade A, Alexandersson E, Levander F. Normalyzer: a tool for rapid evaluation of normalization methods for omics data sets. J Proteome Res. 2014; 13(6):3114–20.

[81] Tyanova S, Temu T, Sinitcyn P, et al. The Perseus computational platform for comprehensive analysis of (prote)omics data. Nat Methods. 2016; 13(9):731–40.

[82] Vizcaíno JA, Csordas A, del-Toro N, et al. 2016 update of the PRIDE database and its related tools. Nucleic Acids Res. 2016; 44(22):11033.

